# DNA double-strand break yield and radiation quality of diagnostic X-rays from 40 to 120 kV: a scale-resolved microdosimetric and track-structure study

**DOI:** 10.64898/2026.08.02.742272

**Authors:** Toshioh Fujibuchi

## Abstract

Reported relative biological effectiveness (RBE) values for low-energy X-rays disagree, assays scoring initial DNA double-strand breaks (DSBs) returning about 1.1 and chromosome-level assays 2 to 4. Whether radiation quality varies within the diagnostic range, and how its comparison with a megavoltage reference depends on target scale, has not been quantified on a tube-potential series. A tungsten-anode tube with 1 mm Be and 2.5 mm Al filtration, with copper added in some cases, was modelled in PHITS for 40 to 200 kV. The spectra were transported into a water phantom in which absorbed dose, lineal-energy densities and cluster size distributions were scored for target diameters of 3 nm to 1 micrometre against a cobalt-60 reference; DSB yields were computed in the electron track-structure mode with the PHITS DNA damage tally. Between 40 and 120 kV the depth-dose ratio changed by a factor of 5.7 and the tube output by a factor of 42, whereas the dose-mean lineal energy varied by 2.5 % at 1 micrometre and 1.2 % at 3 nm against a reproducibility of 0.3 %. Relative to cobalt-60 it was 2.05 times larger at 1 micrometre but only 1.08 times larger at 3 nm, while DSB yields per unit dose were 5 to 7 % higher and constant across the range within the 2 % bound set by the statistics. Tube potential therefore changes the amount and distribution of dose but not its physical quality, and a stated RBE is incomplete without the target scale implied by the endpoint.

## 1. Introduction

Diagnostic radiology accounts for the largest component of population exposure to man-made ionising radiation. Tube potentials in clinical use lie mostly between 40 and 120 kV. In the present system of radiological protection the radiation weighting factor *w*_R_ for photons is unity irrespective of energy [19], so that soft X-rays and ^60^Co γ-rays are taken to carry the same stochastic risk per unit absorbed dose.

Experimental radiobiology does not support this equivalence unambiguously. For mammographic beams, γ-H2AX focus assays give an RBE of 1.4 relative to ^60^Co and micronucleus assays 3–4 at low doses [13], whereas a 53BP1 assay against ^137^Cs gives 1.1 ± 0.2 [14]; comparisons of 25 kV with 200 kV give values mostly between 1 and 2, and below 1 for one micronucleus endpoint [15]. Absorbed dose is a mean over a volume containing many tracks and carries no information on how energy is distributed within them. Microdosimetry supplies that through the lineal energy *y* = ε/*l̅*, the energy imparted to a target of diameter *d* divided by the mean chord length *l̅* = 2*d*/3, and its dose probability density *d*(*y*) [18]; the dose-mean lineal energy *y*_D_ correlates with photon RBE [5,6,7]. On smaller scales, nanodosimetry counts the number ν of ionisations produced in a target of a few nanometres, and *F*_2_ = *P*(ν ≥ 2) is used as a physical surrogate for double-strand break induction.

Much of what follows is already known, and it is worth saying so at the outset. Kellerer showed in 2002 that between 20 and 100 keV — the range that matters for diagnostic radiology — the transition from photoelectric to Compton energy transfer causes a transient decrease in the energy of the released electrons, so that the ionisation density of 200 kVp and 30 kVp beams is similar, and that the low-dose RBE of mammographic against conventional X-rays must therefore be substantially below two [23]. Verhaegen and Reniers found the mean cluster order in 10 nm spheres to be about 35 % higher for mammographic X-rays than for 300 keV electrons but found no significant difference in 2 nm spheres [25]. Hsiao and Stewart obtained an RBE for double-strand break induction of 1.16 for 29 kVp against ^60^Co, against 1.15 measured [26]. Most directly, Tang et al. irradiated endothelial cells at 40 kVp, 220 kVp and 4 MV, simulated all three in Geant4-DNA, and reported few differences between 40 and 220 kVp in either microdosimetric or nanodosimetric quantities, equivalent break yields per gray and per gigabase pair, and similar break complexity [24]. The scale argument is older still: Nikjoo and co-workers concluded in 1991 that effects depending on complex local damage may vary substantially between low-LET radiations while those depending on simple local damage may not [27].

The more recent literature has concentrated on the contrast between kilovoltage and megavoltage [5,6,17,28]. What has not been done is a systematic series. The kilovoltage results above are isolated pairs or triplets of beam qualities, computed in different codes with different geometries, reference radiations and target sizes, so the size of the variation *within* the diagnostic range, and its dependence on the scale at which it is measured, cannot be read off from them. The present work therefore sets out to confirm and to quantify: seven tube potentials from 40 to 200 kV and two copper-filtered variants, one geometry, one transport code, one reference quality, and six target diameters from 3 nm to 1 µm, with the numerical reproducibility of every comparison stated. Absorbed dose, lineal energy, cluster size and explicit strand-break yields are computed from the same simulations, so that the macroscopic and the DNA-scale behaviour of the same beams can be set side by side. The clinical practice of trading tube potential against tube current is conducted entirely in terms of absorbed dose, and the question this series is meant to answer is how much room that practice leaves for a change in radiation quality.

## 2. Materials and Methods

Full details of the tube model, the half-value-layer simulations, the parameter settings, the history budgets and the numerical verification are given in the Supplementary Material, Sections S1–S4.

### 2.1 Transport and beam generation

All calculations used PHITS 3.35 [20] with the EGS5 algorithm (`negs` `=` `1`) for photons, electrons and positrons, and its defaults for electron-impact ionisation, K and L fluorescence and Doppler broadening. A tungsten anode was irradiated by a normally incident electron beam behind a 1 mm Be window and 2.5 mm Al filtration; 0.1 or 0.2 mm Cu was added at 100 kV. Photons emerging at take-off angles of 8–35° were tallied in 0.5 keV bins, using 168 million primary electrons in total across seven tube potentials and four random seeds. The characteristic-line structure of the resulting spectra is resolved: Kα_1_/Kα_2_ = 1.72 at 200 kV against an accepted atomic value near 1.7 (Supplementary Section S1). First half-value layers, obtained by narrow-beam simulation, rise from 1.33 mm Al at 40 kV to 3.63 mm at 120 kV; they lie 6–27 % below the IEC 61267 RQR values, with a shortfall that grows with tube potential (Supplementary Section S2).

### 2.2 Phantom and scored quantities

The tabulated spectra were used as a source on a 20 cm water phantom in which four coaxial discs of 4 cm radius and 5 mm thickness were placed at depths of 0.25, 2, 5 and 10 cm. Absorbed dose and the microdosimetric quantities were scored in these discs, with 2 cm depth used throughout as the reference. ^60^Co γ-rays were modelled as an equal mixture of 1.17 and 1.33 MeV photons in the same geometry.

Lineal-energy probability densities were obtained with the `[T-SED]` tally in its improved form (`model` `=` `1`), which applies an analytical microdosimetric function, derived from track-structure calculations and validated between 3 nm and 1 µm, to the charged-particle information produced by macroscopic transport [1–3]. It is not itself a track-structure simulation. Target diameters of 3, 10 and 25 nm, 100 nm, 377.2 nm and 1 µm were used, spanning the DNA double helix with its hydration shell, the nucleosome, the chromatin fibre and the cell nucleus; 377.2 nm is the domain diameter of the stochastic microdosimetric kinetic model distributed with PHITS [4]. From *f*(*y*) and *d*(*y*) come the frequency and dose means *y*_F_ and *y*_D_, the saturation-corrected *y*\*, and the single-event specific energy *z*_F_ = 4*y*_F_/(ρπ *d*^2^).

With `se-unit = 0` the same tally returns the probability distribution of the number ν of ionisations **and electronic excitations** in the target. This is not quite the ionisation cluster size distribution of the nanodosimetry literature, which counts ionisations alone — the mean energy imparted divided by the mean cluster size gives 22–24 eV rather than the roughly 30 eV per ionisation in water — and the quantities are labelled accordingly. It was scored at 3, 10 and 25 nm on a unit-width mesh, giving *M*_1_, *F*_2_ = *P*(ν ≥ 2) and *F*_3_ = *P*(ν ≥ 3), all conditional on ν ≥ 1.

The electron transport cut-off was 10 keV, as prescribed for `[T-SED]`, which accounts analytically for δ-rays below it; the sensitivity to this choice is examined in Supplementary Section S4. Reproducibility of *y*_D_ is 0.3 % for the X-ray beams and about 0.5 % for ratios referred to ^60^Co (Supplementary Section S3).

### 2.3 Track-structure simulation and DNA damage yields

To test whether the conclusions drawn from these quantities survive a calculation that models strand breakage explicitly, the electron track-structure mode of PHITS was used with the DNA damage estimation code of Matsuya and co-workers [9,21,22]. This required a locally compiled executable; PHITS 3.35 was rebuilt with GNU Fortran 16.1.0 after replacing `usrtally.f90` by the version distributed with the damage code (Supplementary Section S5).

A single electron was started at the centre of a water sphere of 5 mm radius, which exceeds the continuous-slowing-down range of a 1 MeV electron, so the whole track including all secondary electrons is contained and the yields refer to a complete track. The coordinates of all inelastic interactions were written by `[T-Userdefined]` and analysed by the damage code, which superimposes a statistical model of nuclear DNA on the track, converts ionisations and excitations into direct and indirect strand breaks, and classifies double-strand breaks by the number of accompanying lesions within ten base pairs. Yields are reported per unit absorbed dose and per dalton of DNA. Complete tracks were computed for fourteen energies between 2 keV and 1 MeV, between 8 and 68 tracks per energy.

Yields for a beam quality were obtained by weighting the single-track yields by the spectrum of the electrons the photons set in motion. That spectrum, and not the slowing-down fluence, is the correct weight, because a track started at 100 keV already contains its own passage through every lower energy. It was tallied with `[T-Product]` using `output` `=` `atomic` in the same scoring disc, with the electron cut-off lowered to 1 keV. The tally cannot distinguish a δ-ray from an electron released by a photon, so δ-rays were removed afterwards using their spectra measured separately for mono-energetic electrons of the same fourteen energies (Supplementary Section S5).

## 3. Results

### 3.1 Absorbed dose against radiation quality

The macroscopic behaviour of the beams changes strongly with tube potential. The tube output rises as approximately the 3.4 power of the tube potential, a factor of 42 between 40 and 120 kV, and the ratio of dose at 10 cm depth to dose in the entrance disc rises from 0.033 to 0.188, a factor of 5.7 (Supplementary Table S2).

The microdosimetric behaviour does not. Between 40 and 120 kV the dose-mean lineal energy *y*_D_ varies by 2.5 % at the 1 µm target and 1.2 % at 3 nm (Figure 1a, Supplementary Table S3):

**Figure 1.**
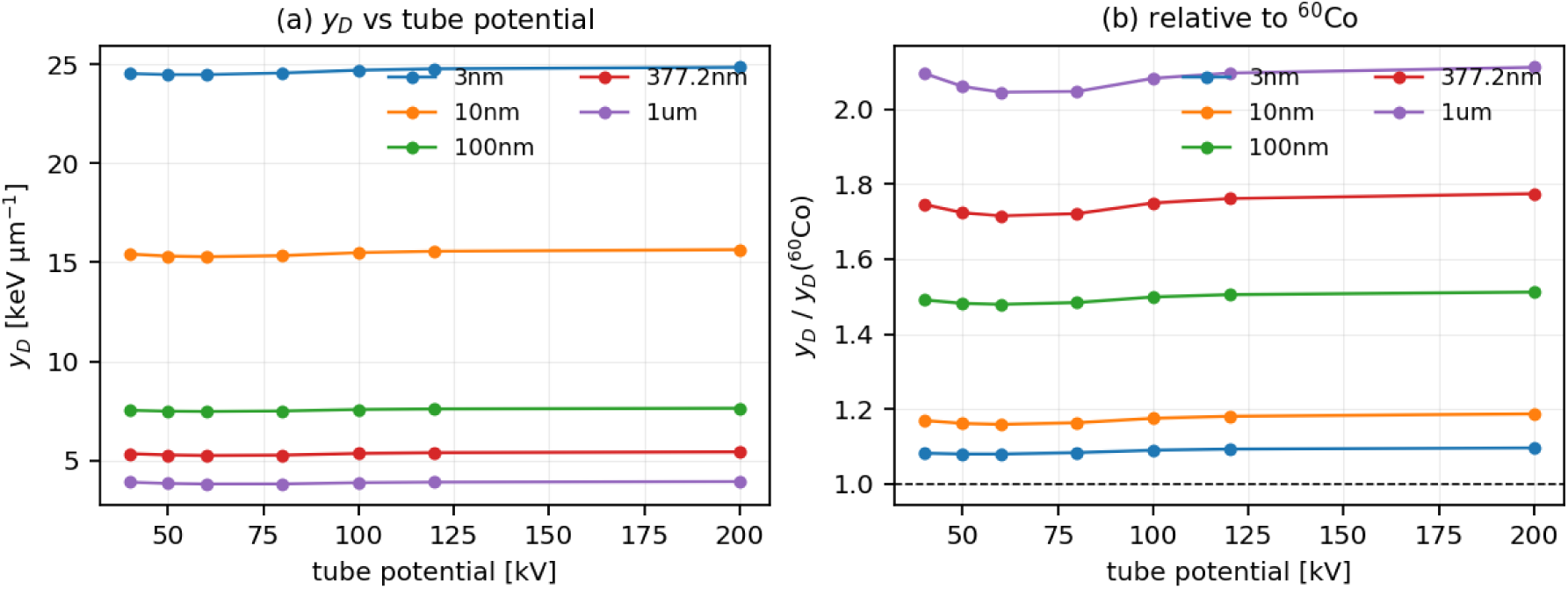
Tube-potential and target-size dependence of the dose-mean lineal energy. (a) *y*_D_ against tube potential for each target diameter. (b) The same values divided by the ^60^Co result. *y*_D_ is nearly independent of tube potential at every target size, whereas the ratio to ^60^Co increases systematically with target size, from about 1.1 at 3 nm to about 2.0 at 1 µm.

| Tube potential [kV] | 40 | 50 | 60 | 80 | 100 | 120 | 200 |
| --- | --- | --- | --- | --- | --- | --- | --- |
| $y_D$ (1 $\mu\text{m}$) [keV $\mu\text{m}^{-1}$] | 3.897 | 3.833 | 3.805 | 3.808 | 3.873 | 3.900 | 3.928 |
| $y_D$ (3 nm) [keV $\mu\text{m}^{-1}$] | 24.53 | 24.48 | 24.48 | 24.56 | 24.71 | 24.77 | 24.85 |

The dependence is not monotonic: *y*_D_ rises towards both ends of the range. Against a reproducibility of 0.3 % the spread is resolved but the position of the minimum is not, the 60 and 80 kV values differing by 0.08 %. Adding 0.1 or 0.2 mm of copper at 100 kV, which raises the mean photon energy by 12 % and 19 %, changes *y*_D_ at 1 µm by 1.1 %. Between 0.25 and 10 cm depth the absorbed dose from a 40 kV beam falls by a factor of thirty while *y*_D_ changes by 4.7 %; at 120 kV the dose falls fivefold and *y*_D_ by 0.6 % (Supplementary Section S6).

### 3.2 Dependence on target size

The comparison with ^60^Co depends strongly on the target diameter (Figure 1b). At 60 kV:

| Target diameter | 3 nm | 10 nm | 25 nm | 100 nm | 377 nm | 1 $\mu\text{m}$ |
| --- | --- | --- | --- | --- | --- | --- |
| $y_D$ ratio (60 kV / $^{60}\text{Co}$) | 1.08 | 1.16 | 1.30 | 1.48 | 1.72 | 2.05 |
| $F_2$ ratio | 1.10 | 1.51 | 1.82 | — | — | — |
| $M_1$ ratio | 1.01 | 1.12 | 1.39 | — | — | — |

At the scale of a cell nucleus the ratio is about two; at the scale of the DNA double helix it is 1.08, and the mean cluster size in a 3 nm target differs by only 1 %. Reducing the target from 1 µm to 3 nm raises *y*_D_ itself by a factor of about six, far exceeding any effect of tube potential.

### 3.3 Why the dependence is weak

For mono-energetic photons *y*_D_ is not monotonic in energy but passes through a minimum of 3.62 keV µm^-1^ near 40 keV and a local maximum of 4.32 keV µm^-1^ near 80 keV before falling towards the megavoltage value (Figure 2a). Below 40 keV the photoelectric effect dominates and the photoelectron carries almost the whole photon energy, so lowering the photon energy lowers the electron energy and raises the stopping power; above 40 keV the growing Compton contribution supplies recoil electrons of only a few keV, which are densely ionising; above about 80 keV those recoils themselves become energetic and sparsely ionising. The mean photon energies of the 40–120 kV spectra, 27.5–51.7 keV, lie entirely inside the basin between about 25 and 60 keV over which *y*_D_ varies by ±5 %.

**Figure 2.**
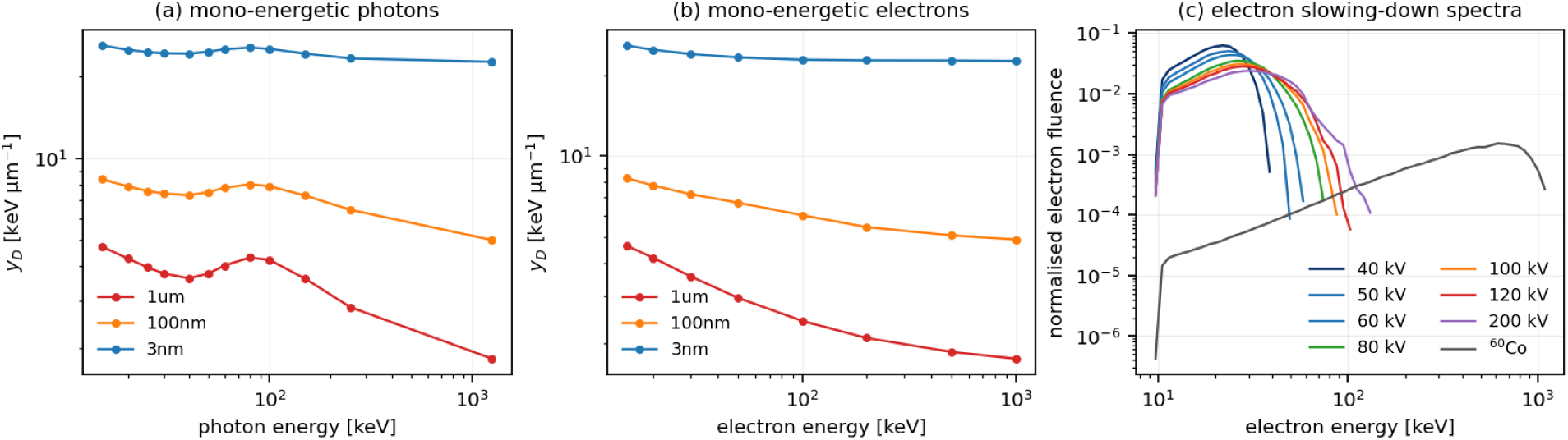
Physical origin of the tube-voltage insensitivity. (a) *y*_D_ for mono-energetic photons, showing a minimum near 40 keV; the mean energies of all diagnostic spectra fall within this shallow basin. (b) *y*_D_ for mono-energetic electrons from 15 keV to 1 MeV. The dependence on electron energy is strong at the 1 µm target and weak at 3 nm, so the near-invariance of the beams reflects compensation within the dose-weighted mixture of secondary electron energies rather than insensitivity. The value at 500 keV, near the mean energy of Compton electrons released by 1.25 MeV photons, coincides with that obtained for the ^60^Co beam. (c) Electron slowing-down spectra in the scoring disc, normalised to unit area. The diagnostic beams produce electron spectra confined below about 120 keV and differing only modestly among themselves, whereas the ^60^Co spectrum extends to 1 MeV.

The compensation should not be mistaken for insensitivity. For mono-energetic electrons *y*_D_ at 1 µm falls from 4.63 to 1.76 keV µm^-1^ between 15 keV and 1 MeV (Figure 2b), and the electron field itself changes appreciably: the mean energy of secondary electrons above the transport cut-off rises from 23.7 keV at 60 kV to 29.4 keV at 120 kV while the fraction below 30 keV falls from 75 % to 57 % (Figure 2c). What is nearly constant is the dose-weighted mixture, because raising the photon energy simultaneously makes the photoelectrons faster, which lowers *y*_D_, and shifts a larger share of the dose to slow Compton recoils, which raises it. At the 3 nm target the same electron-energy change alters *y*_D_ by only 12 %, because the DNA-scale value is dominated by track ends whose ionisation density is set by the last few hundred electronvolts and is therefore common to every photon quality. Neither the shape of the curve nor its consequence is new; Kellerer set out this mechanism from electron spectra in 2002 [23]. What the present series adds is how flat the region is, measured with the reproducibility stated and at target sizes from 3 nm to 1 µm rather than at one scale.

### 3.4 Yield of clustered ionisations per unit absorbed dose

*F*_2_ is a conditional probability and is not by itself the quantity that determines damage per unit dose, because the number of events per unit dose also differs between qualities. The combination needed is the number of targets receiving two or more ionisations per unit dose, which is the cluster dose of Faddegon et al. [29] restricted to ν ≥ 2.

At absorbed dose *D* a target receives *D*/*z*_F_ energy-deposition events, but not all of them can contain an ionisation: *z*_F_ averages over every event that deposits energy, including those below the lowest electronic excitation of liquid water, whereas *F*_2_ is conditioned on at least one ionisation or excitation. Putting the two on a common population introduces a factor \phi = *P*(ε > ε_th_):

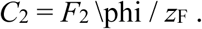

With ε_th_ = 7 eV, the lowest electronic excitation of water, \phi is 0.798 for 40 kV and 0.760 for ^60^Co at 3 nm. The correction differs between qualities and does not cancel from a ratio; omitting it reverses the sign of the result. Relative to ^60^Co (Figure 3, Supplementary Table S7):

**Figure 3.**
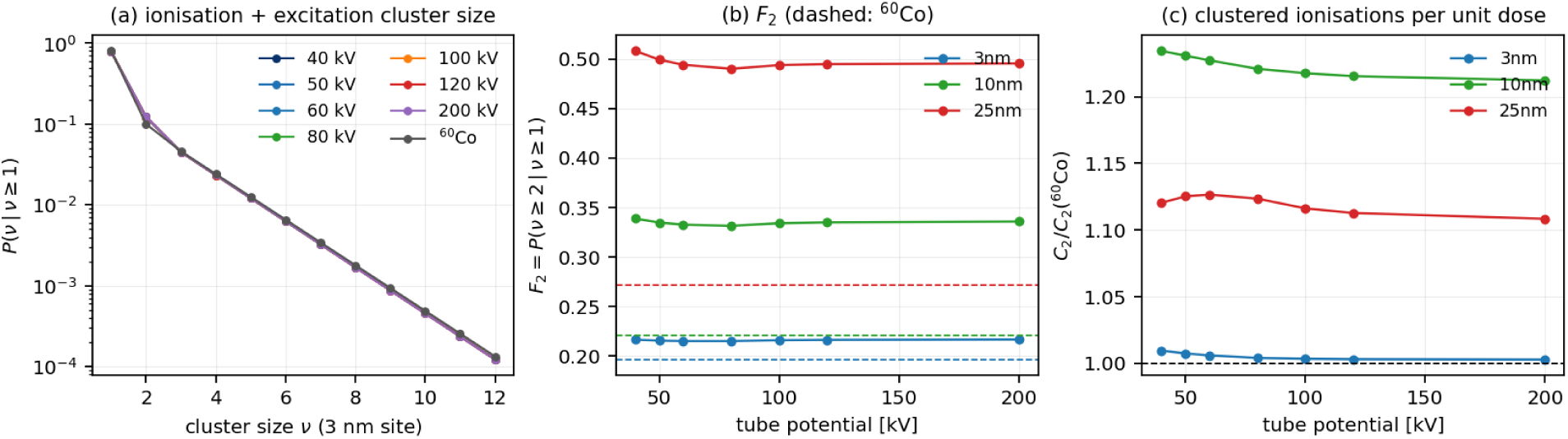
Nanodosimetry. (a) Distributions of the number of ionisations and electronic excitations in a 3 nm target, conditional on at least one event; all beam qualities including ^60^Co nearly coincide. (b) The probability *F*_2_ of two or more such events in a target, against tube potential for three target diameters; dashed lines give the corresponding ^60^Co values. (c) The yield of targets receiving two or more ionisations per unit absorbed dose, *C*_2_ = *F*_2_ \phi/*z*_F_, relative to ^60^Co, where \phi restricts *z*_F_ to the events that can contain an ionisation. Although *F*_2_ per event is higher for the diagnostic beams, they produce fewer events per gray, and at the 3 nm DNA scale the two effects cancel to within 1 % on the central assumption; the threshold and cluster-order sensitivities widen that to about 15 % (Section 3.4).

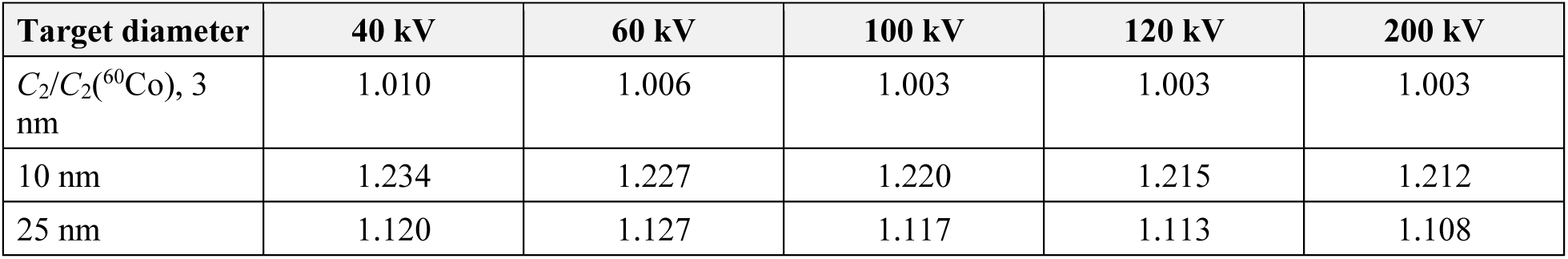

At the 3 nm scale the diagnostic beams and ^60^Co produce the same number of clustered ionisations per unit dose to within 1 %, constant across the tube-potential range to 0.7 %; the enhancement is largest at the 10 nm scale of the nucleosome, about 22 %.

Two sensitivities belong with the 3 nm figure, which is far closer than the model can resolve. Taking ε_th_ as the ionisation potential of water, 10.8 eV, gives 1.039–1.046, and 20 eV gives 1.18–1.19. And a double-strand break requires at least two breaks on opposing strands, so ν ≥ 3 is at least as defensible a surrogate: the corresponding *C*_3_ ratios are 0.886–0.890, because although ^60^Co has the lower probability of exactly two ionisations it has the higher probability at every ν ≥ 3. Together these place the 3 nm ratio between about 0.89 and 1.19 — unity within some 15 % — and it is that bound rather than the central value that the comparison supports. Absolute nanodosimetric quantities carry a further systematic uncertainty of order 30 % from the choice of low-energy cross sections [12], which is expected but not demonstrated to divide out of a ratio between two photon qualities dominated by the same track ends.

### 3.5 DNA damage yields from track-structure simulation

Along a complete electron track the single-strand break yield is almost independent of energy, rising by 3.3 % between 2 keV and 1 MeV, whereas the double-strand break yield falls by 31 %, from 2.082 to 1.431 × 10^-11^ Gy^-1^ Da^-1^ (Figure 4a). Almost all of that change occurs below 40 keV; between 40 keV and 1 MeV the yield varies by 2.7 %, comparable with the standard error of the individual points. The fraction of breaks classified as complex rises from 20 % at 2 keV to 27–29 % above 25 keV and is then constant (Figure 4b). That trend should not be interpreted: the track-structure cut-off used here, 7 eV, is the value of the distributed input, whereas the model was calibrated with 1 eV [22], and repeating the 2 keV point at 1 eV raises the complex fraction to 26.7 % while changing the break yield by 2 %.

**Figure 4.**
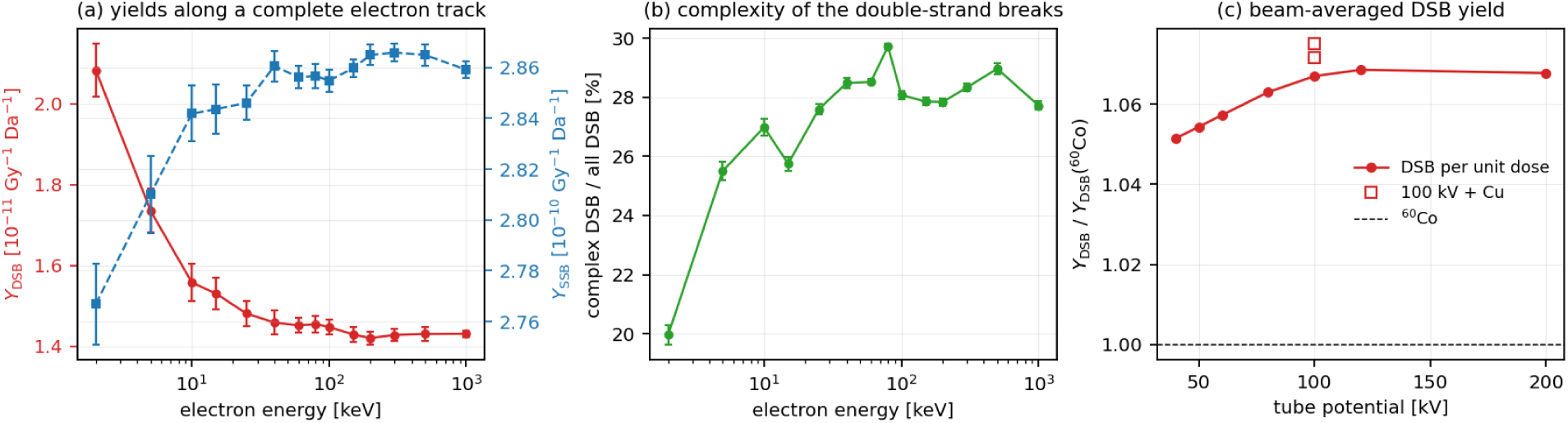
DNA damage yields from track-structure simulation. (a) Yields of single- and double-strand breaks along a complete electron track, per unit absorbed dose and per dalton of DNA, against initial electron energy. The single-strand break yield is almost constant; the double-strand break yield falls by 31 % between 2 keV and 1 MeV, almost all of it below 40 keV. (b) Proportion of double-strand breaks accompanied by at least one further lesion within ten base pairs. The low-energy end of this curve is partly an artefact of the 7 eV track-structure cut-off used here (Section 3.5). (c) Double-strand break yield per unit dose for each beam quality, obtained by weighting (a) with the spectrum of electrons the photons set in motion, and divided by the ^60^Co value. Error bars in (a) and (b) are standard errors of the mean over the simulated tracks; in (c) they are propagated from the simulated energies that carry the spectral weight and are of the order of 1–1.5 %, comparable with the whole vertical range of the panel.

Folding the single-track yields over the spectrum of electrons each beam sets in motion (Figure 4c, Supplementary Table S17) gives (1.505 ± 0.022) × 10^-11^ Gy^-1^ Da^-1^ at 40 kV and (1.530 ± 0.014) × 10^-11^ at 120 kV. The ratio between them is 1.016 ± 0.018: the yield is the same at the two tube potentials within the statistics of the calculation, which bound any difference at about 2 %. The dose-weighted mean energy of the electrons responsible rises only from 25.1 to 30.3 keV over that interval, placing every diagnostic beam on the flat part of Figure 4a. Copper filtration changes the yield by 0.4–0.8 %, again within the statistics. Relative to ^60^Co, whose yield is (1.432 ± 0.007) × 10^-11^ Gy^-1^ Da^-1^, the ratios are 1.051 ± 0.016 at 40 kV and 1.069 ± 0.011 at 120 kV. The complexity is unchanged: 27.3 % of breaks are complex for every diagnostic beam against 28.1 % for ^60^Co, within the bootstrap uncertainty, so no ordering is resolved.

Two limits on the precision belong with these numbers. The fold is dominated by very few simulated energies — 66 % of the weight for 40 kV sits on the 25 keV point and 56 % of the weight for ^60^Co on the 1 MeV point, which has eight tracks — so the quoted uncertainties are governed by those ensembles rather than by the spectra. Removing δ-rays from the production spectra shifts every ratio by 1.0–1.5 % in the same direction for all beams, so the comparison between qualities does not depend on that correction.

Expressed for a nucleus of 6.4 Gbp at 650 Da per base pair the yields are 63 breaks per gray for the diagnostic beams and 60 for ^60^Co, roughly twice what focus assays report; the calculation counts every break whereas an assay counts what it can resolve, and the conversion assumes a DNA content. The argument rests on the ratio, not the absolute level.

## 4. Discussion

### 4.1 Two classes of quantity, two to three orders of magnitude apart

The tube-voltage dependences separate cleanly:

| Quantity | 40 → 120 kV | largest spread over 40–120 kV |
| --- | --- | --- |
| Tube output (energy fluence per incident electron) | × 42 | × 42 |
| Depth-dose ratio, 10 cm / surface | × 5.7 | × 5.7 |
| $y_D$ (1 $\mu\text{m}$) | 0.1 % | 2.5 % |
| $y_D$ (3 nm) | 1.0 % | 1.2 % |
| $C_2$ (3 nm) | 0.7 % | 0.7 % |
| $Y_{\text{DSB}}$, track structure | $1.6 \pm 1.8$ % | $1.6 \pm 1.8$ % |
| Low-dose RBE, microdosimetric kinetic model | 0.5 % | 1.4 % |

Absorbed dose and its distribution are what tube potential and filtration control; radiation quality is not. This is a confirmation rather than a discovery — Tang et al. reached the same conclusion in Geant4-DNA for 40 against 220 kVp with γ-H2AX measurements alongside [24], and Kellerer had predicted it from electron spectra two decades earlier [23] — but it is now quantified across a series, at six target sizes, with the numerical reproducibility stated. The practical consequence is direct: dose reduction achieved by raising the tube potential or hardening the beam is not offset by an increase in initial physical damage per unit dose. Since the ionisation density of the track cannot be lowered by any choice of tube potential or filtration, protection in this energy range depends on reducing dose.

### 4.2 Scale dependence, and what it does and does not explain

The difference from a megavoltage reference is not one number but a function of the scale at which the comparison is made, running from 1.08 at 3 nm to 2.05 at 1 µm. The shape of that dependence is not new either: Verhaegen and Reniers found mammographic X-rays some 35 % above 300 keV electrons at 10 nm but indistinguishable at 2 nm [25], and Nikjoo and co-workers had concluded from cylindrical nanometre targets that endpoints depending on complex local damage may vary appreciably between low-LET radiations while those depending on simple local damage may not [27]. The present series adds the same comparison carried continuously across six diameters for one set of beams, and two estimates of what it implies for double-strand breaks: the clustered-ionisation yield, equal to ^60^Co within 1 % centrally and 15 % on the sensitivities, and the explicit strand-break calculation, 1.051 ± 0.016 at 40 kV.

These two are less independent than they appear. Both are cluster analyses of ionisation and excitation events in homogeneous liquid water, sharing the same cross sections and the same absence of chromatin geometry, hydration shell and explicit chemical stage; the damage model differs in counting strand breaks with fitted coefficients rather than ionisations, not in the underlying track structure. Their agreement is a consistency check, not corroboration by an independent method. That check has been made elsewhere: Hsiao and Stewart obtained 1.16 for 29 kVp against ^60^Co with PENELOPE and a separate damage model, against 1.15 measured [26], and Tang et al. found equivalent yields for 40 and 220 kVp in Geant4-DNA [24]. The present values sit slightly below Hsiao and Stewart’s, as expected, 29 kVp being softer than any beam studied here.

It is tempting to read the scale dependence as an account of why reported RBE values disagree, with DSB assays responding to nanometre clustering and returning values near unity and chromosome or micronucleus assays responding to a micrometre-scale property and returning values near two. These calculations do not support that reading.

The assignment of a target diameter to an assay has no independent criterion, and the table above supplies a continuum of ratios from 1.08 to 2.09, so any reported value in that interval can be matched after the fact. A *y*_D_ ratio is not an RBE: turning one into the other needs a response function, and every response function applied here gives a smaller number than the raw ratio — the stochastic microdosimetric kinetic model gives 1.28–1.31 at its 377 nm domain, and Okamoto et al., fitting a microdosimetric kinetic model to measured survival, obtained an RBE near 1.1 for 200 kV X-rays against megavoltage beams while measuring a *y*_D_ ratio of 1.93 at 1 µm [7]. That is the same pairing used here for the TEPC comparison, and it contradicts a naive reading of the 1 µm ratio by a factor of two. Nor is the experimental record ordered by scale: Habelt and Dörr report micronucleus frequencies giving an RBE between 0.6 and 0.8 for 25 kV against 200 kV alongside values between 1 and 2 for other endpoints in the same study [15]; Guerrero-Carbajal et al. find the initial slope for dicentrics varying by about a factor of five, exceeding the largest ratio available at any target size here, and conclude that their data do not support the view that lower photon energies carry greater risk per unit dose [16]; Lindborg et al. find the α-ratio to follow the *y*_D_ ratio most closely at about 10 nm [17], where the table gives 1.16, even though α is a clonogenic coefficient. Finally, the studies compared differ in reference radiation, dose range, dose rate, cell system and detection threshold as well as in endpoint, and several of the mechanisms documented inside those same papers are individually sufficient: Depuydt et al. attribute their γ-H2AX/micronucleus difference to repair [13], and the values of 3–4 are low-dose limits of a linear-quadratic fit, where the curvature of the ^60^Co response alone will produce α-ratios of that size.

What these results do establish is narrower. The physical quality of the radiation is essentially constant across the diagnostic range at every scale examined; its difference from a megavoltage reference depends on the scale at which the comparison is made, by a factor of two between 3 nm and 1 µm; and a statement of the form “the RBE of diagnostic X-rays is *x*” is therefore incomplete unless the target scale implied by the endpoint is specified. Whether the observed spread among reported values is in fact ordered by that scale is a question these calculations raise and cannot answer.

### 4.3 Implications for radiological protection

Two statements need to be kept apart. Across tube potentials, radiation quality is constant to within a few per cent from 40 to 120 kV, so there is no physical case for making a biological weighting a function of tube potential within the diagnostic range. Across biological scales, the effectiveness relative to ^60^Co is not a single number: about 1.0 judged by double-strand break induction, about 1.3 by the microdosimetric kinetic model with its 377 nm domain, and about 2.0 by *y*_D_ at the 1 µm scale of a cell nucleus. Physics does not select among these; the choice depends on which endpoint carries the risk, which is a radiobiological and epidemiological question.

No change to *w*_R_ is proposed. Such a change would have to be consistent with the reference quality of the underlying epidemiological risk coefficients and would propagate through the system of dose limits. What these results support is the more limited statement that the effectiveness of diagnostic X-rays relative to megavoltage reference qualities is at most a factor of about two, is scale dependent, and does not vary with tube potential.

At diagnostic dose levels the energy deposition is sparse. The single-event specific energy in a 1 µm target is 362–406 mGy for the diagnostic beams against 65.6 mGy for ^60^Co, so at equal dose ^60^Co produces five to six times as many events, each correspondingly smaller. At 1 mGy the mean number of events in a 1 µm volume is 0.003, so such volumes are essentially never hit twice and the specific energy received by one that is hit is fixed by the radiation quality and independent of the macroscopic dose. Reducing dose reduces the number of damaged volumes but not the severity of each event.

### 4.4 Limitations

`[T-SED]` is not a track-structure simulation but an analytical function applied to condensed-history transport, assuming spherical targets in homogeneous water [1–3]; real chromatin is heterogeneous and DNA carries a hydration shell and is wrapped around histones. The explicit strand-break calculation is not subject to that particular objection but has limitations of its own: its DNA model is statistical rather than geometric, its indirect component enters through fitted parameters rather than a chemical stage, and it has been validated for electrons and protons only [9,21,22].

The tube model assumes normal electron incidence, averages over take-off angles of 8–35°, and omits the glass envelope, insulating oil and housing port. The consequence is quantified: half-value layers lie 6–27 % below the IEC 61267 RQR values, and the residual after adding 1.0 mm Al equivalent runs from +7 % at 40 kV to −12 % at 120 kV, so missing inherent filtration accounts for part of the discrepancy but not for its tube-potential dependence. The beams studied are therefore somewhat softer than the corresponding standardised qualities by an amount that grows with tube potential; since *y*_D_ changes by 2.5 % over a factor of 2.7 in half-value layer, a 20 % shift in quality corresponds to well under 1 % in *y*_D_.

Four features of the calculations should be weighed with the results. The half-value-layer simulations used the original PHITS photon transport rather than EGS5, unlike everything else reported here. The copper shell is the outermost layer of the modelled tube, so its 8 keV K fluorescence escapes without the aluminium backing a real filter would carry; it is a negligible part of the dose and does not affect the microdosimetric quantities, but it depresses the first half-value layer of the two copper-filtered beams, and the question whether beams of equal half-value layer produced by different filter materials have the same quality is therefore left open. All production runs used a single random seed, so the reproducibility quoted above is bounded by comparisons that differ in history count and spectral binning rather than measured from seed-to-seed scatter. And the track-structure calculations used a 7 eV electron cut-off where the damage model was calibrated with 1 eV, which affects the complexity curve of Figure 4b though not the beam averages, the dose in every beam being carried by electrons of 20–40 keV where the two settings agree.

Only one anode angle and one field size were studied, the calculations are for water rather than tissue, and no biological experiment accompanies them.

## 4.5 Conclusion

Between 40 and 120 kV the microdosimetric and nanodosimetric character of diagnostic X-rays is essentially constant, although the macrodosimetric character changes by factors of two to forty. This confirms, on a systematic seven-point series in a single framework, what had been inferred from electron spectra [23] and found for isolated pairs of beam qualities in other transport codes [24,25]. Tube potential and added filtration determine how much dose is delivered and where, but not the physical quality of that dose, so that dose reductions achieved by raising the tube potential or hardening the beam are not offset by any increase in initial physical damage per unit dose. The same holds with depth in tissue.

The reason, set out by Kellerer in 2002 [23], is that *y*_D_ has a minimum near 40 keV photon energy, where the photoelectric and Compton contributions to the dose exchange roles. The mean energies of all diagnostic spectra lie within this shallow basin, so changing the tube potential moves the spectrum along an almost flat part of the curve.

The difference from a megavoltage reference depends on the biological target scale. Per unit absorbed dose, diagnostic X-rays and ^60^Co produce the same yield of clustered ionisations at the 3 nm scale of the DNA double helix to within about 15 %, because the higher clustering probability per event of the diagnostic beams is cancelled by their smaller number of events per gray. A track-structure calculation that counts strand breaks explicitly agrees, putting the double-strand break yield per unit dose 5–7 % above ^60^Co and constant across the diagnostic range within its statistics, with no resolved change in the complexity of the breaks. At the 1 µm scale of a cell nucleus, however, *y*_D_ is about twice as large. A statement of the form “the RBE of diagnostic X-rays is *x*” is therefore incomplete unless the target scale implied by the endpoint is specified; whether the spread among reported RBE values is in fact ordered by that scale is not established here, the published measurements differing in reference radiation, dose range, dose rate, cell system and detection threshold as well as in endpoint.

For radiological protection a single effectiveness value suffices across tube potentials. Its magnitude relative to a megavoltage reference is about 1.0 on the basis of double-strand break induction, 1.3 on the basis of the microdosimetric kinetic model domain and 2.0 on a cell-nucleus basis, and the choice between them is a radiobiological rather than a physical one. No change to the radiation weighting factor is proposed.

## Data availability

The PHITS input decks, the analysis scripts and the complete set of numerical tables are provided as Supplementary Material.

