## supplement for "DNA double-strand break yield and radiation quality of diagnostic X-rays from 40 to 120 kV: a scale-resolved microdosimetric and track-structure study"

**Toshioh Fujibuchi**

Division of Medical Quantum Science, Department of Health Sciences, Faculty of Medical Sciences, Kyushu University, 3-1-1 Maidashi, Higashi-ku, Fukuoka 812-8582, Japan

**Short running title:** Radiation quality of X-rays from 40 to 120 kV

**Corresponding author:** Toshioh Fujibuchi, Division of Medical Quantum Science, Department of Health Sciences, Faculty of Medical Sciences, Kyushu University, 3-1-1 Maidashi, Higashi-ku, Fukuoka 812-8582, Japan. Tel: [to be inserted]. E-mail: [to be inserted]. ORCID: [to be inserted].

**Keywords:** microdosimetry; nanodosimetry; DNA double-strand break; diagnostic X-rays; relative biological effectiveness; Monte Carlo simulation

Prepared for submission to the *Journal of Radiation Research* as a Regular Paper.

### S1. Tube model and photon spectra

#### X-ray tube model

The tube was modelled explicitly rather than represented by a semi-empirical spectrum generator, so that the entire study rests on one transport code. A thick tungsten anode of density 19.30 g $cm^{-3}$ occupied the half-space $z < 0$ inside a sphere of radius 0.50 cm, with the focal spot at the origin. A mono-energetic electron pencil beam of kinetic energy equal to the tube potential was directed along $-z$ at normal incidence. Filtration consisted of a 1.0 mm beryllium shell at 0.50–0.60 cm radius and a 2.5 mm aluminium shell at 0.60–0.85 cm, both concentric with the focal spot so that the path length through the filters is independent of emission angle; in a subset of runs a copper shell of 0.1 or 0.2 mm was added outside the aluminium.

Photons were scored in the part of a spherical shell of radius 2.0–2.2 cm lying between $z = 0.30$ and $z = 1.20$ cm, a volume of 2.375 $cm^3$ corresponding to polar angles of 55–82° and hence to X-ray take-off angles of 8–35° with a mean near 20°. Normal electron incidence is an accurate simplification because in a tube with a 20° anode angle the electron beam meets the anode surface at about 20° from its normal, so that the self-filtration path length differs by \cos 20° = 0.94. The approximation makes the problem axially symmetric about $z$, which permits a much larger scoring solid angle and correspondingly better statistical efficiency.

Transport cut-offs were 10 keV for electrons and positrons and 8 keV for photons. Photons below about 12 keV cannot penetrate 2.5 mm of aluminium, so the photon cut-off does not affect the emergent spectrum. Spectra were scored in 0.5 keV bins for tube potentials of 40, 50, 60, 80, 100, 120 and 200 kV, a resolution chosen so that the tungsten $K\alpha_2$ line at 57.98 keV and the $K\alpha_1$ line at 59.32 keV, separated by 1.34 keV, are resolved from one another.

Only a small fraction of the electron energy emerges as photons and only about 21 % of the emitted solid angle is scored, so high statistics are expensive. Each beam was computed as four statistically independent runs with different random seeds and the results averaged, which keeps all four physical cores of the machine working on the slowest beam. Histories were scaled inversely with tube output so that all beams end with comparable counts per bin, from 60 million primary electrons at 40 kV to 8 million at 120 kV and 168 million in total. The uncertainty of the averaged spectrum was obtained both by propagating the per-bin errors and, independently, from the scatter between the four seeds; the two agree to better than one percentage point, for example 9.9 % against 9.9 % at 40 kV and 12.5 % against 13.0 % at 60 kV.

**Photon spectra**

The spectra generated by the tube model are shown in Figure S1 and summarised in Table S1. The 2.5 mm Al filtration removes all photons below about 13–15 keV. Tungsten characteristic lines appear on the bremsstrahlung continuum only for tube potentials above the K-shell ionisation threshold of 69.5 keV and are absent at 40, 50 and 60 kV.

At the 0.5 keV binning the four K lines are individually resolved (Figure S1c): $K\alpha_2$ at 57.98 keV, $K\alpha_1$ at 59.32 keV, $K\beta_{1,3}$ at 67.24 keV and $K\beta_2$ at 69.07 keV. Their relative intensities are atomic constants and therefore provide a test of the tube model that is independent of the transport of the continuum. After subtraction of a local continuum baseline the calculated $K\alpha_1/K\alpha_2$ intensity ratio is 1.92 at 100 kV, 1.79 at 120 kV and 1.72 at 200 kV, converging on the accepted value of about 1.7 as the statistics improve, and the $K\beta/K\alpha$ ratio at 200 kV is 0.26, within the accepted range of 0.23–0.28.

Fluence-weighted mean photon energies are 27.5 keV at 40 kV, 31.2 keV at 50 kV, 34.6 keV at

60 kV, 41.0 keV at 80 kV, 46.6 keV at 100 kV, 51.7 keV at 120 kV and 66.0 keV at 200 kV. The tube output, taken as the energy fluence per incident electron, rises by a factor of 42 between 40 and 120 kV, corresponding to a power-law exponent of 3.4 in tube potential; over the narrower interval 60–120 kV the exponent is 3.0. Both values lie within the range of 2.5–3.5 usually observed at fixed filtration.

Copper hardens the 100 kV beam as expected. The mean photon energy rises from 46.6 keV with 2.5 mm Al alone to 52.0 keV with 0.1 mm Cu and 55.5 keV with 0.2 mm Cu, while the output falls to 74 % and 59 % of its value with aluminium alone. A small copper K-fluorescence component at 8.0 keV is present in the copper-filtered spectra, as it is in practice.

#### S2. Half-value layer

##### Half-value layer

Beam quality was characterised by the first and second half-value layer (HVL) in aluminium, obtained by direct simulation. Each spectrum was used as an on-axis pencil beam incident on an aluminium slab of thickness 0 to 10 mm, with an air detector of 0.5 cm radius placed 1 m behind the slab so that photons scattered in the aluminium diverge away from it, which is the good-geometry condition of the HVL definition. Electron transport was suppressed by raising the electron cut-off above the maximum photon energy, so that `[t-deposit]` reverts to the kerma approximation and the scored quantity is the collision air kerma in which HVL is defined.

The response of this detector was verified by irradiating it with mono-energetic photons and dividing the scored kerma by the photon energy fluence, which returns the mass energy-absorption coefficient of air. The result follows the expected behaviour of $(\mu_{en}/\rho)_{air}$, falling from 1.36 $cm^2\ g^{-1}$ at 15 keV through a broad minimum of 0.023 $cm^2\ g^{-1}$ near 80–100 keV and rising to 0.027 $cm^2\ g^{-}$

$^{1}$ at 200 keV, and agrees with tabulated values to within a few per cent over the whole range.

**Half-value layer**

The simulated first half-value layers in aluminium rise from 1.33 mm at 40 kV to 3.63 mm at 120 kV and 6.02 mm at 200 kV (Table S15, Figure S8a). Homogeneity coefficients $HVL_1/HVL_2$ fall from 0.85 at 40 kV to 0.62 at 120 kV, as expected for progressively broader spectra.

These values lie 6–27 % below the HVLs specified for the IEC 61267 RQR series at the same tube potential, and the shortfall increases monotonically with tube potential. The model contains only a beryllium window and the added aluminium, whereas a real tube also has a glass envelope, insulating oil and a housing port. Because the transmission curve of each beam was measured over a range of absorber thicknesses, the effect of additional filtration can be evaluated without further simulation: the HVL of the same beam pre-hardened by $f$ mm of aluminium is the extra thickness required to halve the kerma that survives the first $f$ mm. Adding 1.0 mm Al equivalent gives 2.32 mm against 2.19 mm at 60 kV, 3.08 mm against 3.01 mm at 80 kV and 3.88 mm against 3.97 mm at 100 kV.

That agreement does not extend across the range. Against the same 1.0 mm column the residual runs from +7 % at 40 kV through +2 % at 80 kV to −12 % at 120 kV, and a constant added filtration cannot remove a residual that changes sign. Missing inherent filtration therefore accounts for part of the discrepancy but not for its tube-potential dependence, which points instead to a deficient high-energy bremsstrahlung continuum in the anode model. The beams studied here are consequently somewhat softer than the corresponding standardised qualities, by an amount that grows with tube potential; the consequences for the microdosimetric results are considered in Section 4.4 of the main text.

The offset does not affect the quantities of interest here. Over the clinical range the HVL varies by a factor of 2.7 while $y_D$ varies by 2.5 % (Section 3.1 of the main text), so a 20 % shift in beam quality corresponds to a shift in $y_D$ of well under 1 %. Plotted against HVL rather than tube potential, $y_D$ is flat from 1.3 to 6 mm Al (Figure S8b).

Copper filtration raises the simulated HVL of the 100 kV beam from 3.11 mm to 3.42 mm with 0.1 mm Cu and 4.99 mm with 0.2 mm Cu, and the two copper-filtered points fall close to the aluminium-filtered line in Figure S8b. That comparison is, however, not a fair one in the present geometry. The copper shell is the outermost layer of the modelled tube, so its 8.0 keV K fluorescence leaves without being absorbed; it amounts to well under 1 % of the fluence but, being far softer than the rest of the beam, it dominates the first half-value layer. Its signature is visible in Table S15: the first HVL rises by 1.6 mm Al for the second 0.1 mm of copper against 0.3 mm for the first, and the homogeneity coefficient falls to 0.55, whereas copper filtration should narrow the spectrum and raise it. In practice a copper filter is backed by aluminium precisely to remove this line. The copper-filtered $y_D$ values are unaffected, the fluorescence carrying a negligible share of the dose, but their abscissa in Figure S8b is not, and the question whether beams of equal half-value layer produced by different filter materials have the same microdosimetric quality is therefore left open by these calculations.

#### S3. Statistics, computational details and numerical verification

##### Computational details

Tube spectra were generated with 60, 30, 16, 12, 10, 8 and 9 million primary electrons at 40, 50, 60, 80, 100, 120 and 200 kV respectively and 11–12 million for each copper-filtered case, each split over four random seeds. The median relative uncertainty per 0.5 keV bin is 10 % at 40 kV and 12–15 % elsewhere; bins containing a characteristic line have smaller errors, 5 % for $K\alpha_1$ at 100 kV, because the line concentrates counts into a single bin. Phantom calculations used 150 000 source photons for each beam quality, 25 000 for the mono-energetic photon scan, 2 000 for the mono-energetic electron scan, in which every history deposits its whole energy inside the scoring volume, and 40 000 for $^{60}$Co, whose long electron tracks make each history expensive.

All phantom and spectrum runs were executed in freshly created directories with `istdev = 1`. With `istdev = -1` the code resumes from any tally files left in the working directory and merges the earlier statistics into the new run without warning. The track-structure runs of Section S5 are the one exception: they use `istdev = -2` and restart deliberately, which is how the damage code accumulates tracks.

##### Numerical verification

The microdosimetric function accounts analytically for δ-rays below the transport cut-off, and the code documentation specifies an electron cut-off of 10 or 100 keV. Because most Compton recoil electrons produced by 40–120 kV X-rays are born below 10 keV, the sensitivity to this setting was examined by repeating calculations with cut-offs of 1, 10 and 100 keV.

##### Comparison with measurement

Table S14 compares the calculated $y_D$ at the 1 μm target with published TEPC measurements [7]. The absolute values obtained here are 13–20 % lower, 3.93 against 4.51 keV $\mu m^{-1}$ for 200 kV X-rays and 1.86 against 2.34 keV $\mu m^{-1}$ for $^{60}$Co, whereas the ratio between the two qualities is 2.11 here against 1.93 measured, a difference of 9 %.

#### S4. Sensitivity to the electron transport cut-off

##### Sensitivity to the electron transport cut-off

Repeating the calculations with electron cut-offs of 1, 10 and 100 keV gives the results in Table S12 and Figure S7. For $y_D$ at the 1 µm target:

| Cut-off | 40 kV | 120 kV | difference |
|---|---|---|---|
| 1 keV | 3.306 | 3.239 | −2.0 % |
| 10 keV | 3.896 | 3.910 | +0.4 % |
| 100 keV | 3.911 | 3.908 | −0.1 % |

The 10 and 100 keV settings, both sanctioned by the code documentation, agree to within 3 % at every target size, the largest discrepancy being 2.7 % at the 10 nm target and 0.4 % or better at 1 µm. A 1 keV cut-off gives systematically lower $y_D$, by 15 % at 1 µm and 7 % at 3 nm, as expected because it transports δ-rays that the microdosimetric function already accounts for analytically, so that their contribution is in effect counted twice and the lineal-energy spectrum is redistributed.

The comparison between tube potentials is preserved under every choice of cut-off: the difference in $y_D$ between 40 and 120 kV is at most 2.1 % whichever setting is used and is of the same order as the numerical reproducibility. All results reported elsewhere use the 10 keV setting.

#### S5. Track-structure simulation: build, settings and the electron production spectrum

##### Track-structure simulation and DNA damage yields

The lineal-energy and cluster-size quantities of Section S3 and 2.6 describe the physical pattern of energy deposition but say nothing about DNA itself. To test whether the conclusions drawn from them survive a calculation that models strand breakage explicitly, the electron track-structure mode of PHITS was used together with the DNA damage estimation code of Matsuya and co-workers [9, 21, 22], which is distributed with the code as a user tally.

Track-structure mode follows every elastic and inelastic interaction of an electron individually instead of grouping them into condensed-history steps, down to a cut-off of 7 eV. Using it requires a locally compiled executable, because the user tally that writes the interaction coordinates is supplied as Fortran source. PHITS 3.35 was therefore rebuilt from source with GNU Fortran 16.1.0 after replacing `usrtally.f90` by the version distributed with the damage code. One patch was needed: `SPLIT` has become an intrinsic in Fortran 2023, and the compiler resolved a call to the code's own sixteen-argument routine of that name to the intrinsic instead, which was fixed by declaring it `external`.

A single electron was started at the centre of a water sphere of 5 mm radius, which exceeds the continuous-slowing-down range of a 1 MeV electron, so that the whole track including all secondary electrons is contained and the calculation refers to a complete track rather than to a segment of one. The coordinates and types of all inelastic interactions were written by `[T-Userdefined]` and analysed by the damage code, which superimposes a model of the cell nucleus on the track, converts ionisations and excitations into direct and indirect strand breaks, and classifies the double-strand breaks by the number of accompanying lesions within ten base pairs. Yields are reported per unit absorbed dose and per dalton of DNA. The distributed settings were used throughout: proliferating cells in logarithmic phase, complex-damage classification enabled, and 3.4 nm — ten base pairs — as the interaction distance defining complexity. The cell-cycle setting rescales all yields by the mean DNA content of the nucleus and therefore does not affect any ratio reported here.

Complete tracks were computed for fourteen initial energies between 2 keV and 1 MeV. Because the interaction file grows to roughly 0.1 MB per keV of incident energy, each energy was run as a sequence of restarts, the interaction file being analysed and deleted after each. Between 8 and 60 tracks were accumulated per energy, giving a standard error of the mean double-strand break yield of 1–3 %.

Yields for a beam quality were obtained by weighting the single-track yields by the spectrum of the electrons the photons set in motion. That spectrum, and not the slowing-down fluence, is the correct weight, because a track started at 100 keV already contains its own passage through every lower energy. It was tallied with `[T-Product]` using `output = atomic` in the same scoring disc at 2 cm depth, with the electron transport cut-off lowered to 1 keV so that the low-energy Compton electrons of the softer beams are resolved. The tally cannot distinguish a δ-ray from an electron released by a photon, so the δ-rays were removed afterwards: their spectra were measured separately for mono-energetic electrons of the same fourteen energies, and starting from the highest energy bin the δ-rays expected from the electrons already identified as photon-produced were subtracted. The correction is small — a 10 keV electron releases 0.03 δ-rays above 1 keV over its whole track — and results with and without it are compared in Section 3.5 of the main text.

#### S6. Additional results

##### Absorbed dose

Central-axis depth-dose distributions are shown in Figure S2. Normalised to the entrance disc, the dose at 10 cm depth is 0.033 at 40 kV, 0.061 at 50 kV, 0.081 at 60 kV, 0.124 at 80 kV, 0.158 at 100 kV, 0.188 at 120 kV, 0.244 at 200 kV and 0.829 for $^{60}$Co (Table S2). Within the diagnostic range alone this ratio varies by a factor of 5.7. Taken with the factor of 42 in tube output, tube potential is the dominant determinant of the dose distribution.

##### Lineal-energy distributions

Dose probability densities $y\,d(y)$ at 2 cm depth are shown in Figure S3. At a target diameter of 1 μm the curves for all tube potentials from 40 to 200 kV are almost superimposed and only $^{60}$Co is clearly displaced towards lower $y$. At 3 nm the whole distribution moves one to two orders of magnitude higher in $y$, and the separation between the diagnostic beams and $^{60}$Co becomes markedly smaller than at 1 μm.

**Ionisation cluster size distributions**

In a 3 nm target the mean cluster size $M_1$ is 1.411 at both 40 and 120 kV and the probability of two or more ionisations $F_2$ is 0.2166 and 0.2165 respectively, so that the two beams are indistinguishable (Table S6, Figure 3). Relative to $^{60}$Co the $F_2$ ratio is 1.10 at 3 nm, 1.53 at 10 nm and 1.87 at 25 nm, again increasing with target size, while the $M_1$ ratio at 3 nm is 1.01.

**Dependence on depth**

$y_D$ at the 1 μm target is almost independent of depth (Table S13). At 40 kV it changes from 3.968 keV μm$^{-1}$ at 0.25 cm to 3.780 keV μm$^{-1}$ at 10 cm, a fall of 4.7 %; at 60 kV from 3.867 to 3.735 keV μm$^{-1}$, or 3.4 %; and at 120 kV from 3.909 to 3.886 keV μm$^{-1}$, or 0.6 %. Over the same depth range the absorbed dose falls by a factor of 30 at 40 kV and 5 at 120 kV. Beam hardening by preferential absorption of the softer components and beam softening by the growing scattered component evidently compensate almost exactly, the residual being a slight softening.

**Estimated relative biological effectiveness**

The low-dose limit of the stochastic microdosimetric kinetic model, referred to $^{60}$Co, is 1.295 at 40 kV, 1.287 at 50 kV, 1.284 at 60 kV, 1.286 at 80 kV, 1.297 at 100 kV, 1.302 at 120 kV and 1.307 at 200 kV (Table S8). The diagnostic range therefore lies between 1.28 and 1.30, a variation of 1.4 %, again with a shallow minimum near 60–80 kV. Copper filtration changes the value to

1.299 with 0.1 mm and 1.306 with 0.2 mm.

The model parameters were adjusted for tissue reactions rather than for the stochastic effects relevant to protection, and the model contains no description of repair. These values should be read as an index of the difference in initial physical damage on the scale of the model domain, 377 nm, and not as a risk weighting factor.

**Event statistics at diagnostic dose levels**

The single-event specific energy $z_F$ in a 1 µm target falls from 406 mGy at 40 kV to 362 mGy at 120 kV, against 65.6 mGy for $^{60}$Co (Table S9). For equal absorbed dose $^{60}$Co therefore delivers five to six times as many events, each correspondingly smaller.

At 1 mGy the mean number of events in a 1 µm volume is 0.002–0.003 for the diagnostic beams and 0.015 for $^{60}$Co, so that under Poisson statistics 99.7 % of such volumes, or 98.5 % for $^{60}$Co, receive no energy at all. Even at 100 mGy the mean is only 0.25–0.28 events per 1 µm volume for the diagnostic beams.

**Effect of copper filtration**

Adding 0.1 or 0.2 mm of copper to the 100 kV beam raises the mean photon energy from 46.6 keV to 52.0 and 55.5 keV and reduces the tube output to 74 % and 59 % of its value with aluminium alone. The effect on radiation quality is small: $y_D$ at the 1 µm target is 3.873, 3.879 and 3.914 keV µm$^{-1}$ for 2.5 mm Al, +0.1 mm Cu and +0.2 mm Cu, a spread of 1.1 %, and at the 3 nm target 24.71, 24.78 and 24.88 keV µm$^{-1}$ (Figure S6). The yield of clustered ionisations per unit dose at 3 nm, relative to $^{60}$Co, is 1.003, 1.002 and 1.001, and the double-strand break yield from the track-structure calculation rises from 1.067 ± 0.012 to 1.072 ± 0.011 and 1.075 ± 0.011 times the $^{60}$Co value, a change within the statistics.

##### Track structure at diagnostic dose levels

The single-event specific energy $z_F$ in a 1 μm target is 362–406 mGy for the diagnostic beams and 65.6 mGy for $^{60}$Co, so that at equal absorbed dose $^{60}$Co produces five to six times as many events, each correspondingly smaller. At 1 mGy, a realistic organ dose for a radiographic examination, the mean number of events in a 1 μm volume is 0.003 for the diagnostic beams and 0.015 for $^{60}$Co, so that 99.7 % and 98.5 % of such volumes respectively receive no energy at all. The specific energy received by a volume that is hit is fixed by the radiation quality and independent of the macroscopic dose: lowering the dose reduces the number of volumes hit but not the severity of a hit.

Two consequences follow. Because volumes are essentially never hit twice, the physical damage per unit dose cannot depend on dose or dose rate in this regime, and any dose-rate effect that is observed must arise from repair kinetics rather than from a change in the pattern of energy deposition. The absence of intertrack interaction is also the microdosimetric basis for expecting the dose–effect relation to be linear at diagnostic dose levels, though this is an argument about initial damage and not a demonstration concerning cancer risk.

#### S7. Supplementary figures

**Figure S1. Photon spectra emerging from the modelled tungsten-anode tube.** (a) Photon fluence per unit energy and (b) energy fluence per unit energy, each normalised to unit area, for tube potentials of 40–200 kV with 1 mm Be and 2.5 mm Al filtration, scored over take-off angles of 8–35°, in 0.5 keV bins. Vertical grey lines mark the tungsten K-line energies. (c) The K-line region on a linear scale, showing that the binning resolves $K\alpha_2$ (57.98 keV) from $K\alpha_1$ (59.32 keV) and $K\beta_{1,3}$ (67.24 keV) from $K\beta_2$ (69.07 keV).

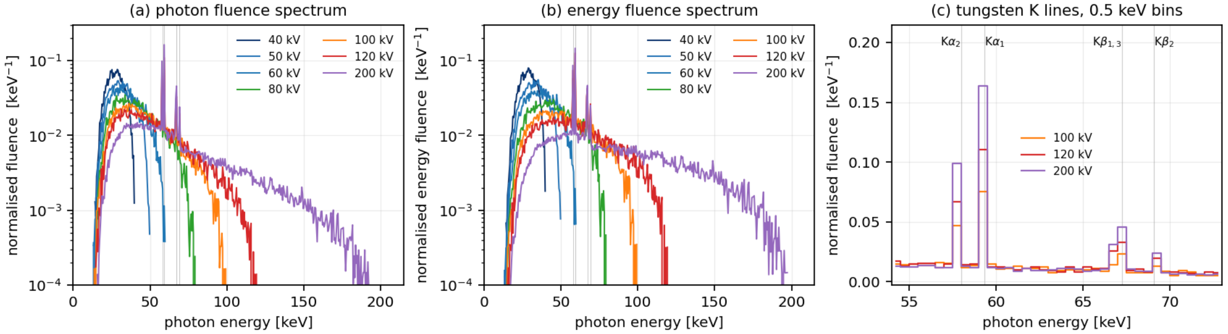


**Figure S2. Central-axis depth-dose distributions in the water phantom**, normalised to the dose in the entrance disc, on (a) logarithmic and (b) linear scales. The curves are volume averages over the 4 cm radius of the scoring discs rather than true central-axis values.

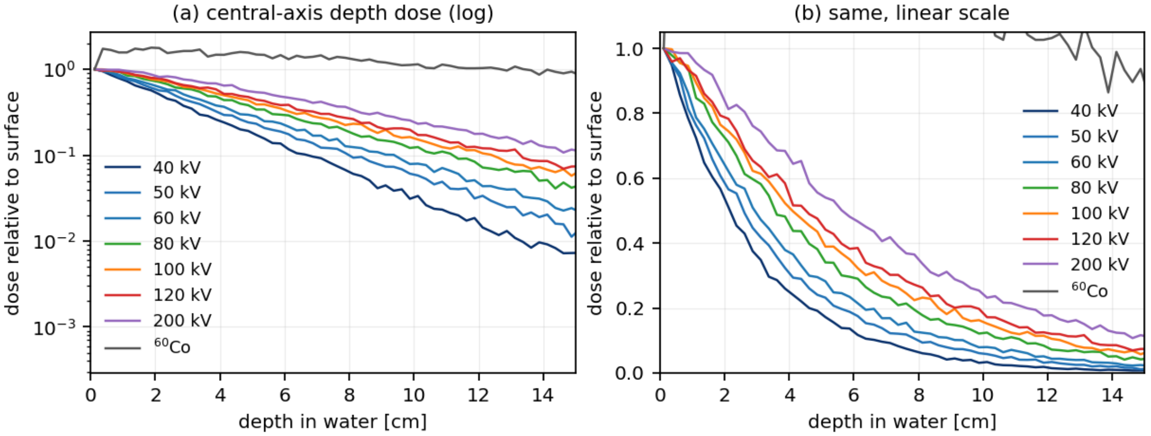


**Figure S3. Dose probability densities of lineal energy, plotted as $y\,d(y)$ against log $y$**, at 2 cm depth for (a) a 1 μm target and (b) a 3 nm target. The diagnostic beams are almost superimposed; only $^{60}$Co is clearly displaced towards lower $y$, and the displacement is much smaller for the 3 nm target.

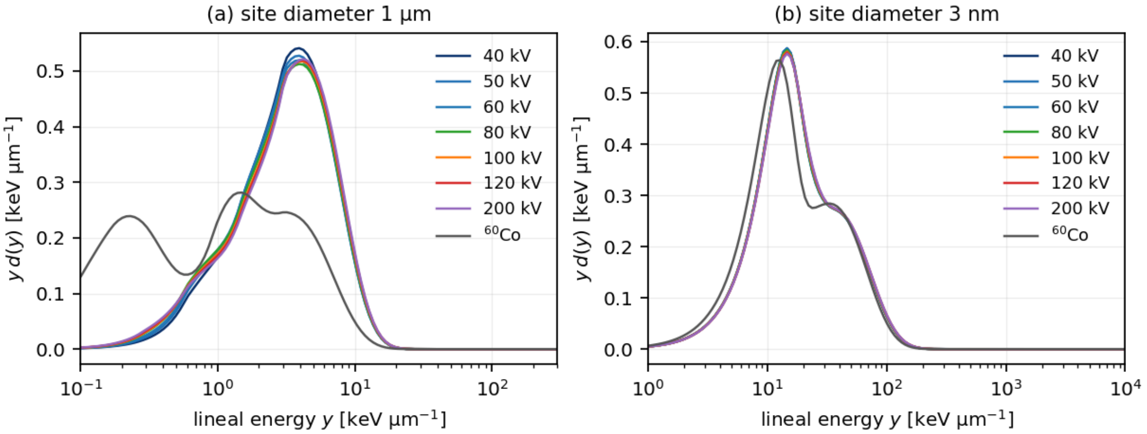


**Figure S4. Dependence of (a) the dose-mean lineal energy $y_D$ and (b) the frequency-mean lineal energy $y_F$ on target diameter** for all beam qualities.

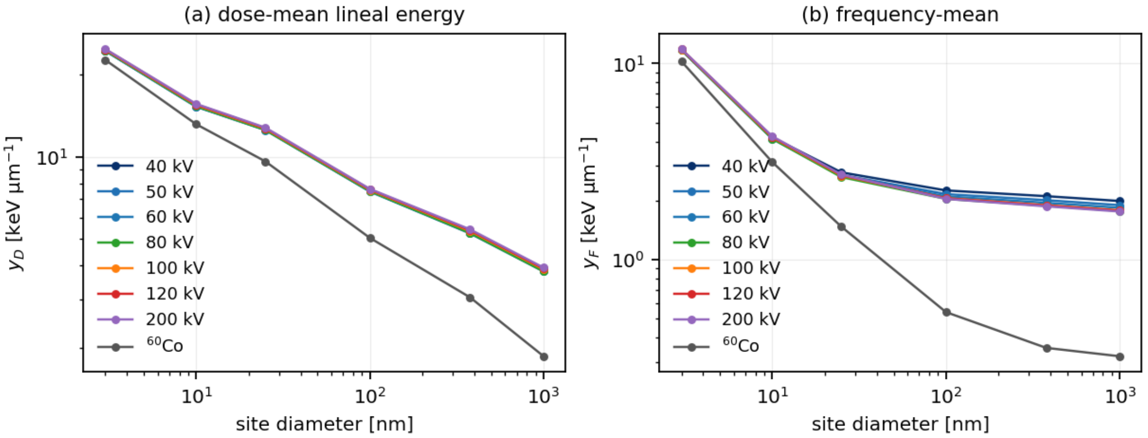


**Figure S5. (a) Low-dose limit of the RBE** predicted by the stochastic microdosimetric kinetic model relative to $^{60}$Co, and **(b) the depth dependence of $y_D$** at the 1 μm target.

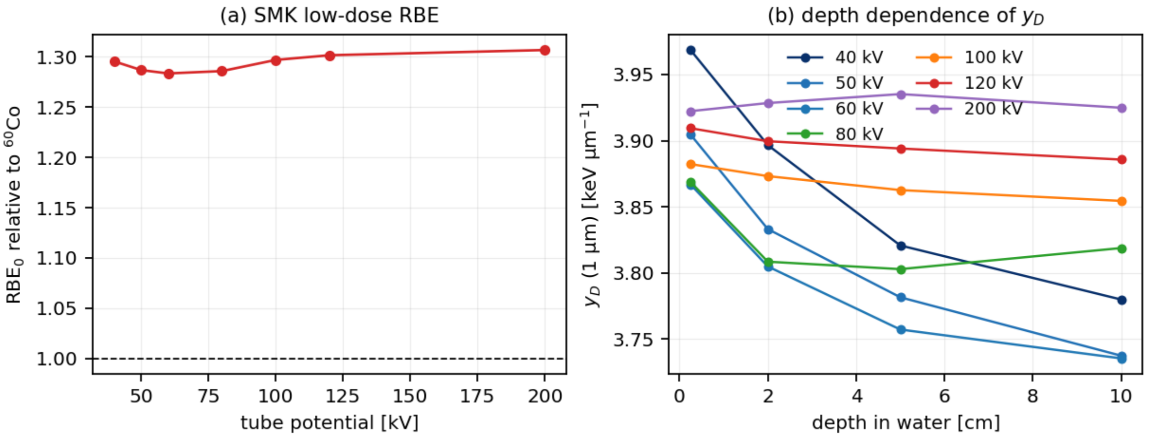

**Figure S6. Effect of added copper filtration at 100 kV.** (a) Normalised photon fluence spectra with 2.5 mm Al alone and with 0.1 or 0.2 mm Cu added. (b) The resulting $y_D$ as a function of target diameter.

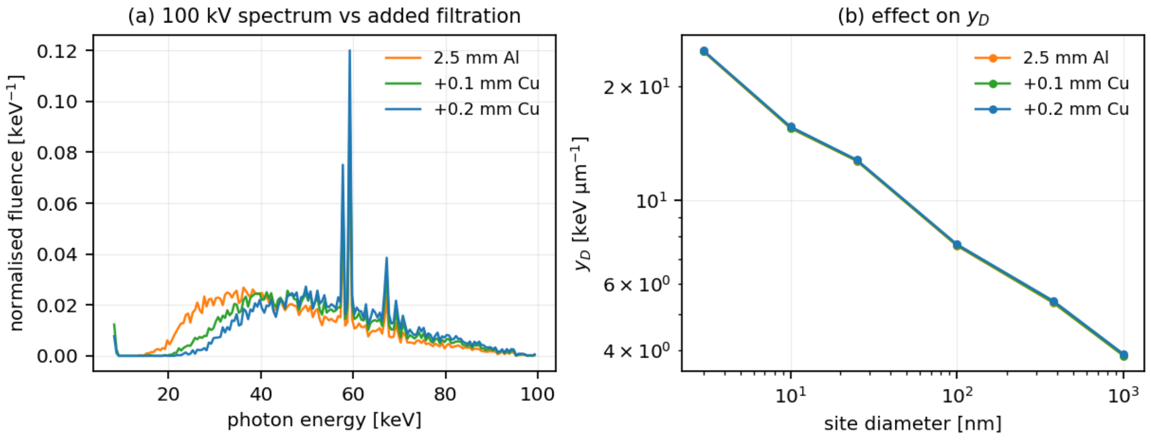


**Figure S7. Sensitivity of $y_D$ to the electron transport cut-off** (1, 10 and 100 keV) for two tube potentials. The 10 and 100 keV settings agree closely; a 1 keV cut-off, which double-counts δ-rays already included analytically in the microdosimetric function, gives systematically lower values.

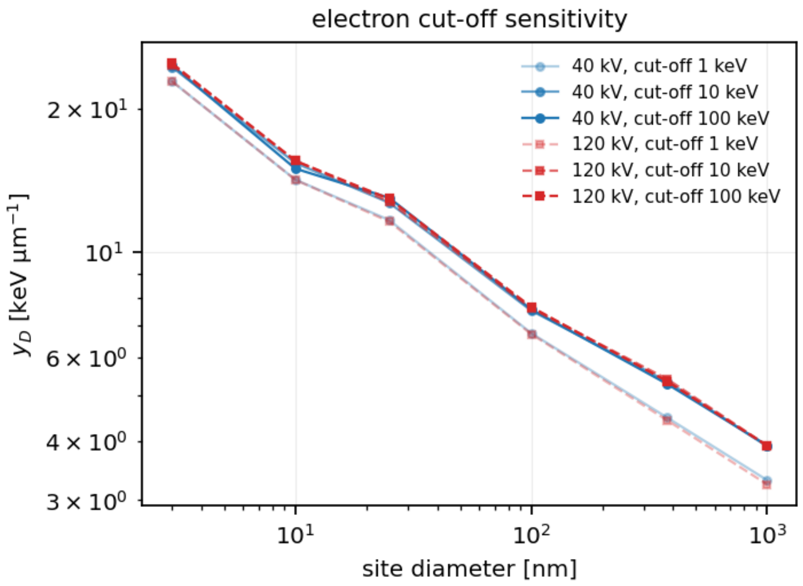


**Figure S8. Beam quality expressed as half-value layer.** (a) First HVL in aluminium against tube potential, from narrow-beam air-kerma transmission curves; the copper-filtered 100 kV beams are shown as squares. (b) $y_D$ at the 1 μm (red, left axis) and 3 nm (blue, right axis) targets plotted against first HVL. The abscissa of the copper-filtered points is not reliable, because the copper shell is the outermost layer of the modelled tube and its 8 keV K fluorescence escapes unfiltered, which depresses the first half-value layer (Section S2); their $y_D$ values are unaffected. What the figure shows is that $y_D$ depends only weakly on beam quality over the whole range.

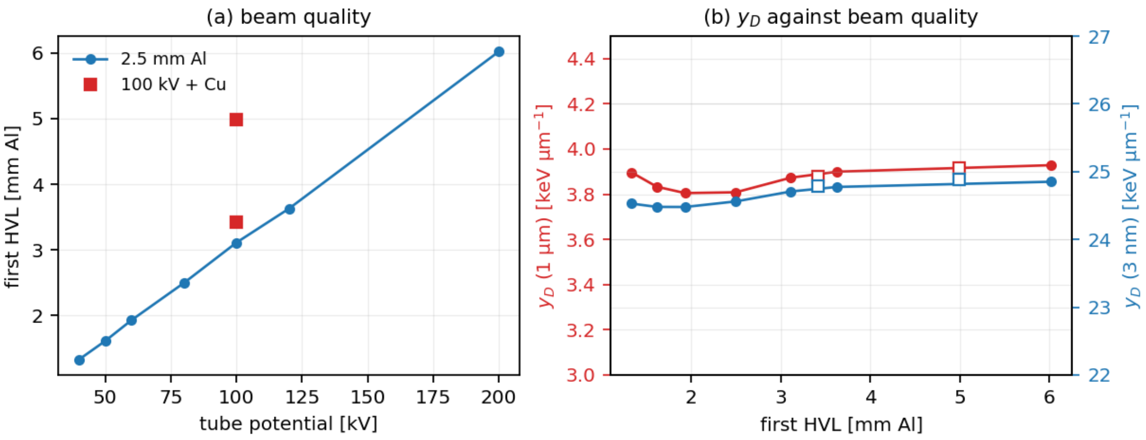


#### S8. Supplementary tables

The complete numerical results follow; the same values are in `results/tables.txt` in the data archive.

##### Table S1 Photon spectra leaving the tube (2.5 mm Al total filtration)

| beam | E_mean [keV] | E_mean,Psi [keV] | E_min [keV] | E_max [keV] | rel. energy fluence per electron |
|---|---|---|---|---|---|
| 40 kV | 27.5 | 28.4 | 14.0 | 40.0 | 1.00 |
| 50 kV | 31.2 | 32.8 | 13.0 | 50.0 | 2.58 |
| 60 kV | 34.6 | 37.1 | 13.0 | 60.0 | 5.15 |
| 80 kV | 41.0 | 45.2 | 14.5 | 79.5 | 12.86 |
| 100 kV | 46.6 | 52.4 | 13.5 | 99.5 | 25.74 |
| 120 kV | 51.7 | 59.0 | 14.5 | 119.5 | 42.10 |
| 200 kV | 66.0 | 80.9 | 15.5 | 197.5 | 135.72 |
| 100 kV +0.1 mm Cu | 52.0 | 56.7 | 8.0 | 99.5 | 18.99 |
| 100 kV +0.2 mm Cu | 55.5 | 59.5 | 8.0 | 99.5 | 15.17 |

##### Table S2 Macrodosimetry: absorbed dose in the scoring discs

| beam | 0.25 cm | 2.00 cm | 5.00 cm | 10.00 cm | D(10)/D(0.25) |
|---|---|---|---|---|---|
| 40 kV | 9.1558e-05 | 5.0674e-05 | 1.7079e-05 | 3.0301e-06 | 0.0331 |
| 50 kV | 7.8573e-05 | 4.7212e-05 | 1.9008e-05 | 4.7805e-06 | 0.0608 |
| 60 kV | 6.9859e-05 | 4.5945e-05 | 2.0627e-05 | 5.6572e-06 | 0.0810 |
| 80 kV | 5.8839e-05 | 4.3188e-05 | 2.1462e-05 | 7.2977e-06 | 0.1240 |
| 100 kV | 5.3920e-05 | 4.0759e-05 | 2.2256e-05 | 8.4910e-06 | 0.1575 |
| 120 kV | 4.9956e-05 | 4.0198e-05 | 2.3389e-05 | 9.4103e-06 | 0.1884 |
| 200 kV | 4.6225e-05 | 3.9038e-05 | 2.5470e-05 | 1.1284e-05 | 0.2441 |
| Co-60 | 3.8525e-04 | 5.0035e-04 | 4.2961e-04 | 3.1924e-04 | 0.8287 |

#### Table S3  Dose-mean lineal energy y_D [keV/um] at 2 cm depth

| beam | 3nm | 10nm | 25nm | 100nm | 377.2nm | 1um |
|---|---|---|---|---|---|---|
| 40 kV | 24.528 | 15.410 | 12.704 | 7.528 | 5.336 | 3.897 |
| 50 kV | 24.479 | 15.310 | 12.591 | 7.482 | 5.269 | 3.833 |
| 60 kV | 24.478 | 15.279 | 12.544 | 7.468 | 5.244 | 3.805 |
| 80 kV | 24.560 | 15.334 | 12.563 | 7.494 | 5.262 | 3.808 |
| 100 kV | 24.707 | 15.488 | 12.690 | 7.568 | 5.348 | 3.873 |
| 120 kV | 24.774 | 15.558 | 12.744 | 7.599 | 5.385 | 3.900 |
| 200 kV | 24.851 | 15.642 | 12.805 | 7.633 | 5.425 | 3.928 |
| 100 kV +0.1Cu | 24.782 | 15.541 | 12.711 | 7.592 | 5.367 | 3.879 |
| 100 kV +0.2Cu | 24.880 | 15.639 | 12.784 | 7.637 | 5.417 | 3.914 |
| Co-60 | 22.666 | 13.179 | 9.633 | 5.049 | 3.057 | 1.860 |

#### Table S4  y_D relative to Co-60

| beam | 3nm | 10nm | 25nm | 100nm | 377.2nm | 1um |
|---|---|---|---|---|---|---|
| 40 kV | 1.082 | 1.169 | 1.319 | 1.491 | 1.745 | 2.094 |
| 50 kV | 1.080 | 1.162 | 1.307 | 1.482 | 1.724 | 2.060 |
| 60 kV | 1.080 | 1.159 | 1.302 | 1.479 | 1.715 | 2.045 |
| 80 kV | 1.084 | 1.164 | 1.304 | 1.484 | 1.721 | 2.047 |
| 100 kV | 1.090 | 1.175 | 1.317 | 1.499 | 1.749 | 2.082 |
| 120 kV | 1.093 | 1.181 | 1.323 | 1.505 | 1.761 | 2.096 |
| 200 kV | 1.096 | 1.187 | 1.329 | 1.512 | 1.774 | 2.111 |
| 100 kV +0.1Cu | 1.093 | 1.179 | 1.319 | 1.504 | 1.756 | 2.085 |
| 100 kV +0.2Cu | 1.098 | 1.187 | 1.327 | 1.513 | 1.772 | 2.104 |
| Co-60 | 1.000 | 1.000 | 1.000 | 1.000 | 1.000 | 1.000 |

#### Table S5  Frequency-mean lineal energy y_F [keV/um]

| beam | 3nm | 10nm | 25nm | 100nm | 377.2nm | 1um |
|---|---|---|---|---|---|---|
| 40 kV | 11.749 | 4.200 | 2.784 | 2.252 | 2.106 | 1.989 |
| 50 kV | 11.715 | 4.148 | 2.702 | 2.158 | 2.009 | 1.895 |
| 60 kV | 11.704 | 4.126 | 2.659 | 2.100 | 1.948 | 1.836 |
| 80 kV | 11.726 | 4.135 | 2.634 | 2.041 | 1.882 | 1.772 |
| 100 kV | 11.791 | 4.195 | 2.679 | 2.057 | 1.894 | 1.783 |
| 120 kV | 11.819 | 4.221 | 2.695 | 2.053 | 1.887 | 1.776 |
| 200 kV | 11.847 | 4.251 | 2.712 | 2.039 | 1.870 | 1.759 |
| 100 kV +0.1Cu | 11.814 | 4.205 | 2.659 | 2.009 | 1.841 | 1.731 |
| 100 kV +0.2Cu | 11.853 | 4.240 | 2.677 | 2.000 | 1.828 | 1.719 |
| Co-60 | 10.249 | 3.140 | 1.471 | 0.540 | 0.355 | 0.321 |

#### Table S6  Cluster statistics for ionisations AND electronic excitations

#### (conditional on nu >= 1; not directly comparable with published ICSDs,

#### which count ionisations alone)

| beam | site | M1 | F2 | F3 |
|---|---|---|---|---|
| 40 kV | 3nm | 1.411 | 0.2166 | 0.0936 |
| 40 kV | 10nm | 1.926 | 0.3389 | 0.1601 |
| 40 kV | 25nm | 2.842 | 0.5081 | 0.2955 |
| 50 kV | 3nm | 1.410 | 0.2157 | 0.0936 |
| 50 kV | 10nm | 1.916 | 0.3349 | 0.1578 |
| 50 kV | 25nm | 2.803 | 0.4992 | 0.2885 |
| 60 kV | 3nm | 1.410 | 0.2153 | 0.0936 |
| 60 kV | 10nm | 1.911 | 0.3327 | 0.1567 |
| 60 kV | 25nm | 2.783 | 0.4941 | 0.2845 |
| 80 kV | 3nm | 1.410 | 0.2153 | 0.0936 |
| 80 kV | 10nm | 1.909 | 0.3316 | 0.1563 |
| 80 kV | 25nm | 2.769 | 0.4901 | 0.2819 |
| 100 kV | 3nm | 1.411 | 0.2161 | 0.0937 |
| 100 kV | 10nm | 1.916 | 0.3341 | 0.1580 |
| 100 kV | 25nm | 2.789 | 0.4937 | 0.2853 |
| 120 kV | 3nm | 1.411 | 0.2165 | 0.0938 |
| 120 kV | 10nm | 1.919 | 0.3350 | 0.1587 |
| 120 kV | 25nm | 2.796 | 0.4948 | 0.2864 |
| 200 kV | 3nm | 1.412 | 0.2169 | 0.0939 |
| 200 kV | 10nm | 1.922 | 0.3359 | 0.1594 |
| 200 kV | 25nm | 2.802 | 0.4955 | 0.2874 |
| 100 kV +0.1Cu | 3nm | 1.411 | 0.2162 | 0.0938 |
| 100 kV +0.1Cu | 10nm | 1.915 | 0.3333 | 0.1578 |
| 100 kV +0.1Cu | 25nm | 2.779 | 0.4904 | 0.2832 |
| 100 kV +0.2Cu | 3nm | 1.411 | 0.2166 | 0.0939 |
| 100 kV +0.2Cu | 10nm | 1.918 | 0.3343 | 0.1586 |
| 100 kV +0.2Cu | 25nm | 2.786 | 0.4913 | 0.2843 |

| | | | | |
|---|---|---|---|---|
| Co-60 | 3nm | 1.399 | 0.1966 | 0.0965 |
| Co-60 | 10nm | 1.710 | 0.2209 | 0.1117 |
| Co-60 | 25nm | 1.998 | 0.2716 | 0.1241 |

#### Table S7  F2 and M1 relative to Co-60

| beam | site | M1 ratio | F2 ratio |
|---|---|---|---|
| 40 kV | 3nm | 1.009 | 1.102 |
| 40 kV | 10nm | 1.127 | 1.534 |
| 40 kV | 25nm | 1.422 | 1.871 |
| 50 kV | 3nm | 1.008 | 1.097 |
| 50 kV | 10nm | 1.120 | 1.516 |
| 50 kV | 25nm | 1.403 | 1.838 |
| 60 kV | 3nm | 1.008 | 1.095 |
| 60 kV | 10nm | 1.117 | 1.506 |
| 60 kV | 25nm | 1.393 | 1.819 |
| 80 kV | 3nm | 1.008 | 1.095 |
| 80 kV | 10nm | 1.116 | 1.501 |
| 80 kV | 25nm | 1.386 | 1.805 |
| 100 kV | 3nm | 1.009 | 1.099 |
| 100 kV | 10nm | 1.120 | 1.513 |
| 100 kV | 25nm | 1.396 | 1.818 |
| 120 kV | 3nm | 1.009 | 1.101 |
| 120 kV | 10nm | 1.122 | 1.517 |
| 120 kV | 25nm | 1.400 | 1.822 |
| 200 kV | 3nm | 1.009 | 1.103 |
| 200 kV | 10nm | 1.124 | 1.521 |
| 200 kV | 25nm | 1.403 | 1.825 |
| 100 kV +0.1Cu | 3nm | 1.009 | 1.100 |
| 100 kV +0.1Cu | 10nm | 1.120 | 1.509 |
| 100 kV +0.1Cu | 25nm | 1.391 | 1.806 |
| 100 kV +0.2Cu | 3nm | 1.009 | 1.102 |
| 100 kV +0.2Cu | 10nm | 1.122 | 1.514 |
| 100 kV +0.2Cu | 25nm | 1.395 | 1.809 |
| Co-60 | 3nm | 1.000 | 1.000 |
| Co-60 | 10nm | 1.000 | 1.000 |
| Co-60 | 25nm | 1.000 | 1.000 |

#### Table S7b  Yield of targets receiving two or more ionisations per unit dose

**C2 = F2 * phi / z_F  [per target per Gy]; only ratios are meaningful.**

**phi = P(eps > threshold) puts F2, which is conditioned on nu >= 1, and**

**z_F, which averages over every energy-deposition event, on the same**

**population.  Columns give the three thresholds of the sensitivity test;**

**'raw' is F2/z_F without the correction.  C3 uses nu >= 3 instead.**

| beam | site | z_F [Gy] | F2 | phi(7eV) | C2 raw | C2 7eV | C2 10.8 | C2 20eV | C3 7eV |
|---|---|---|---|---|---|---|---|---|---|
| 40 kV | 3nm | 266269.1439 | 0.2166 | 0.7976 | 0.961 | 1.010 | 1.046 | 1.190 | 0.889 |
| 40 kV | 10nm | 8567.0384 | 0.3389 | 0.7750 | 1.147 | 1.234 | 1.324 | 1.644 | 1.153 |
| 40 kV | 25nm | 908.6734 | 0.5081 | 0.8154 | 0.988 | 1.120 | 1.247 | 1.681 | 1.426 |
| 50 kV | 3nm | 265494.1788 | 0.2157 | 0.7969 | 0.960 | 1.007 | 1.043 | 1.184 | 0.890 |
| 50 kV | 10nm | 8459.9024 | 0.3349 | 0.7722 | 1.148 | 1.231 | 1.316 | 1.623 | 1.147 |
| 50 kV | 25nm | 881.6972 | 0.4992 | 0.8088 | 1.001 | 1.125 | 1.245 | 1.659 | 1.423 |
| 60 kV | 3nm | 265256.7078 | 0.2153 | 0.7966 | 0.959 | 1.006 | 1.041 | 1.181 | 0.890 |
| 60 kV | 10nm | 8416.3506 | 0.3327 | 0.7710 | 1.146 | 1.227 | 1.311 | 1.612 | 1.143 |
| 60 kV | 25nm | 867.6453 | 0.4941 | 0.8051 | 1.007 | 1.127 | 1.242 | 1.644 | 1.420 |
| 80 kV | 3nm | 265764.9454 | 0.2153 | 0.7966 | 0.957 | 1.004 | 1.039 | 1.179 | 0.889 |
| 80 kV | 10nm | 8434.7141 | 0.3316 | 0.7713 | 1.140 | 1.221 | 1.306 | 1.605 | 1.138 |
| 80 kV | 25nm | 859.7198 | 0.4901 | 0.8021 | 1.008 | 1.124 | 1.235 | 1.626 | 1.414 |
| 100 kV | 3nm | 267237.0281 | 0.2161 | 0.7975 | 0.956 | 1.003 | 1.039 | 1.183 | 0.886 |
| 100 kV | 10nm | 8556.1766 | 0.3341 | 0.7743 | 1.133 | 1.218 | 1.307 | 1.617 | 1.139 |
| 100 kV | 25nm | 874.1988 | 0.4937 | 0.8043 | 0.998 | 1.116 | 1.230 | 1.627 | 1.412 |
| 120 kV | 3nm | 267857.2505 | 0.2165 | 0.7979 | 0.955 | 1.003 | 1.039 | 1.184 | 0.885 |
| 120 kV | 10nm | 8609.3684 | 0.3350 | 0.7756 | 1.129 | 1.215 | 1.307 | 1.620 | 1.138 |
| 120 kV | 25nm | 879.5048 | 0.4948 | 0.8050 | 0.994 | 1.113 | 1.227 | 1.625 | 1.410 |
| 200 kV | 3nm | 268502.7345 | 0.2169 | 0.7981 | 0.954 | 1.003 | 1.039 | 1.185 | 0.884 |
| 200 kV | 10nm | 8670.9501 | 0.3359 | 0.7772 | 1.123 | 1.212 | 1.306 | 1.624 | 1.138 |
| 200 kV | 25nm | 884.9877 | 0.4955 | 0.8056 | 0.990 | 1.108 | 1.223 | 1.622 | 1.407 |
| 100 kV +0.1Cu | 3nm | 267742.0434 | 0.2162 | 0.7976 | 0.954 | 1.002 | 1.038 | 1.182 | 0.885 |
| 100 kV +0.1Cu | 10nm | 8576.9913 | 0.3333 | 0.7747 | 1.127 | 1.212 | 1.302 | 1.611 | 1.135 |

| | | | | | | | | | |
|---|---|---|---|---|---|---|---|---|---|
| 100 kV +0.1Cu | 25nm | 867.8872 | 0.4904 | 0.8018 | 0.999 | 1.113 | 1.224 | 1.612 | 1.407 |
| 100 kV +0.2Cu | 3nm | 268624.0673 | 0.2166 | 0.7981 | 0.953 | 1.001 | 1.037 | 1.183 | 0.884 |
| 100 kV +0.2Cu | 10nm | 8649.1247 | 0.3343 | 0.7765 | 1.121 | 1.208 | 1.301 | 1.615 | 1.134 |
| 100 kV +0.2Cu | 25nm | 873.7496 | 0.4913 | 0.8023 | 0.994 | 1.109 | 1.219 | 1.608 | 1.404 |
| Co-60 | 3nm | 232281.3355 | 0.1966 | 0.7595 | 1.000 | 1.000 | 1.000 | 1.000 | 1.000 |
| Co-60 | 10nm | 6405.6460 | 0.2209 | 0.7202 | 1.000 | 1.000 | 1.000 | 1.000 | 1.000 |
| Co-60 | 25nm | 480.0267 | 0.2716 | 0.7193 | 1.000 | 1.000 | 1.000 | 1.000 | 1.000 |

#### Table S8 SMK model (formulation and parameters as shipped with PHITS,

#### Sato & Furusawa 2012; domain diameter 377.2 nm, y0 = 100 keV/um)

#### z*(Co-60) computed here = 4.362 Gy (value shipped with PHITS: 3.512 Gy)

#### RBE below uses the value computed here, so RBE(Co-60) = 1 exactly.

| beam | y* [keV/um] | z* [Gy] | RBE(D->0) |
|---|---|---|---|
| 40 kV | 5.302 | 7.601 | 1.295 |
| 50 kV | 5.237 | 7.507 | 1.287 |
| 60 kV | 5.211 | 7.470 | 1.284 |
| 80 kV | 5.229 | 7.496 | 1.286 |
| 100 kV | 5.314 | 7.618 | 1.297 |
| 120 kV | 5.349 | 7.669 | 1.302 |
| 200 kV | 5.389 | 7.726 | 1.307 |
| 100 kV +0.1Cu | 5.332 | 7.644 | 1.299 |
| 100 kV +0.2Cu | 5.381 | 7.715 | 1.306 |
| Co-60 | 3.043 | 4.362 | 1.000 |

#### Table S9 Track structure at diagnostic dose levels (1 um site)

| beam | z_F [mGy] | n(1 mGy) | n(10 mGy) | n(100 mGy) | P(0 hit\|1 mGy) |
|---|---|---|---|---|---|
| 40 kV | 405.7827 | 0.002 | 0.025 | 0.246 | 0.9975 |
| 50 kV | 386.5669 | 0.003 | 0.026 | 0.259 | 0.9974 |
| 60 kV | 374.4944 | 0.003 | 0.027 | 0.267 | 0.9973 |
| 80 kV | 361.4004 | 0.003 | 0.028 | 0.277 | 0.9972 |
| 100 kV | 363.6332 | 0.003 | 0.028 | 0.275 | 0.9973 |
| 120 kV | 362.3441 | 0.003 | 0.028 | 0.276 | 0.9972 |
| 200 kV | 358.8215 | 0.003 | 0.028 | 0.279 | 0.9972 |
| 100 kV +0.1Cu | 353.1378 | 0.003 | 0.028 | 0.283 | 0.9972 |
| 100 kV +0.2Cu | 350.5898 | 0.003 | 0.029 | 0.285 | 0.9972 |
| Co-60 | 65.5657 | 0.015 | 0.153 | 1.525 | 0.9849 |

#### Table S10 Mono-energetic photons: y_D [keV/um]

| photon energy [keV] | 3nm | 10nm | 25nm | 100nm | 377.2nm | 1um |
|---|---|---|---|---|---|---|
| 15 | 25.990 | 17.169 | 14.305 | 8.372 | 6.358 | 4.722 |
| 20 | 25.076 | 16.175 | 13.422 | 7.877 | 5.780 | 4.276 |
| 25 | 24.587 | 15.528 | 12.828 | 7.580 | 5.411 | 3.968 |
| 30 | 24.356 | 15.160 | 12.471 | 7.416 | 5.191 | 3.772 |
| 40 | 24.270 | 14.956 | 12.206 | 7.315 | 5.036 | 3.618 |
| 50 | 24.677 | 15.370 | 12.517 | 7.512 | 5.250 | 3.771 |
| 60 | 25.201 | 15.950 | 13.023 | 7.791 | 5.585 | 4.033 |
| 80 | 25.569 | 16.490 | 13.554 | 8.024 | 5.909 | 4.315 |
| 100 | 25.269 | 16.212 | 13.361 | 7.893 | 5.773 | 4.227 |
| 150 | 24.273 | 15.014 | 12.204 | 7.290 | 5.021 | 3.605 |
| 250 | 23.316 | 13.886 | 10.860 | 6.468 | 4.131 | 2.834 |
| 1250 | 22.647 | 13.165 | 9.600 | 5.005 | 3.025 | 1.831 |

#### Table S11 Mono-energetic electrons: y_D [keV/um]

| electron energy [keV] | 3nm | 10nm | 25nm | 100nm | 377.2nm | 1um |
|---|---|---|---|---|---|---|
| 15 | 25.787 | 16.972 | 14.133 | 8.267 | 6.238 | 4.633 |
| 20 | 24.867 | 15.947 | 13.219 | 7.763 | 5.645 | 4.172 |
| 30 | 23.976 | 14.722 | 12.065 | 7.196 | 4.923 | 3.555 |
| 50 | 23.287 | 13.821 | 11.009 | 6.695 | 4.245 | 2.957 |
| 100 | 22.842 | 13.327 | 10.154 | 6.013 | 3.666 | 2.426 |
| 200 | 22.712 | 13.161 | 9.758 | 5.427 | 3.329 | 2.095 |
| 500 | 22.682 | 13.155 | 9.619 | 5.056 | 3.062 | 1.860 |
| 1000 | 22.600 | 13.144 | 9.535 | 4.882 | 2.936 | 1.755 |

#### Table S12 Sensitivity to the electron transport cut-off

| case | 3nm | 10nm | 25nm | 100nm | 377.2nm | 1um |
|---|---|---|---|---|---|---|
| kv040_c001 | 22.883 | 14.207 | 11.679 | 6.746 | 4.477 | 3.306 |
| kv040_c010 | 24.524 | 15.407 | 12.702 | 7.526 | 5.334 | 3.896 |
| kv040_c100 | 24.750 | 14.991 | 13.008 | 7.569 | 5.272 | 3.911 |
| kv120_c001 | 22.909 | 14.192 | 11.590 | 6.715 | 4.419 | 3.239 |
| kv120_c010 | 24.796 | 15.583 | 12.765 | 7.609 | 5.398 | 3.910 |
| kv120_c100 | 24.981 | 15.559 | 12.949 | 7.662 | 5.339 | 3.908 |

#### Table S13 Depth dependence of y_D at the 1 um site [keV/um]

| beam | 0.25 cm | 2.00 cm | 5.00 cm | 10.00 cm |
|---|---|---|---|---|
| 40 kV | 3.968 | 3.897 | 3.821 | 3.780 |
| 50 kV | 3.905 | 3.833 | 3.782 | 3.737 |
| 60 kV | 3.867 | 3.805 | 3.757 | 3.735 |
| 80 kV | 3.869 | 3.808 | 3.803 | 3.819 |
| 100 kV | 3.882 | 3.873 | 3.863 | 3.854 |
| 120 kV | 3.909 | 3.900 | 3.894 | 3.886 |
| 200 kV | 3.922 | 3.928 | 3.935 | 3.925 |
| Co-60 | 1.854 | 1.860 | 1.865 | 1.876 |

#### Table S14 Comparison with published TEPC measurements (1 um site)

#### Okamoto et al., J Radiat Res 52:75 (2011)

| beam | this work | measured | ratio |
|---|---|---|---|
| 200 kV | 3.93 | 4.51 | 0.87 |
| Co-60 | 1.86 | 2.34 | 0.80 |

y_D(200 kV)/y_D(Co-60): this work 2.11 measured 1.93
y_D(40 kV)/y_D(Co-60) : this work 2.09

#### Table S15 Half-value layer in aluminium (narrow-beam air kerma)

#### 'inherent' columns give the HVL the same beam would have if the

#### model also contained the glass/oil/housing filtration of a real tube

| beam | HVL1 | HVL2 | h | +0.5mm | +1.0mm | +1.5mm |
|---|---|---|---|---|---|---|
| 40 kV | 1.33 | 1.57 | 0.85 | 1.46 | 1.52 | 1.61 |
| 50 kV | 1.61 | 2.05 | 0.79 | 1.81 | 1.90 | 2.02 |
| 60 kV | 1.93 | 2.51 | 0.77 | 2.16 | 2.32 | 2.39 |
| 80 kV | 2.50 | 3.42 | 0.73 | 2.82 | 3.08 | 3.16 |
| 100 kV | 3.11 | 4.40 | 0.71 | 3.50 | 3.88 | 3.96 |
| 120 kV | 3.63 | 5.87 | 0.62 | 4.03 | 4.40 | 4.74 |
| 200 kV | 6.02 | - | - | 6.53 | 6.92 | 7.31 |
| 100 kV +0.1Cu | 3.42 | 6.21 | 0.55 | 5.21 | 5.43 | 5.62 |
| 100 kV +0.2Cu | 4.99 | - | - | 6.43 | 6.57 | 6.70 |

#### Table S16 DNA damage along a complete electron track (track structure)

#### Y_SSB, Y_DSB per Gy per Da; complex = (DSB+ and DSB++) / all DSB

| E [keV] | tracks | depe/E | Y_SSB | err% | Y_DSB | err% | complex% | DSB/Gy/cell |
|---|---|---|---|---|---|---|---|---|
| 2 | 60 | 1.000 | 2.7669e-10 | 0.6 | 2.0823e-11 | 3.1 | 20.0 | 86.6 |
| 5 | 60 | 1.000 | 2.8101e-10 | 0.5 | 1.7353e-11 | 3.1 | 25.5 | 72.2 |
| 10 | 50 | 1.000 | 2.8419e-10 | 0.4 | 1.5576e-11 | 3.0 | 27.0 | 64.8 |
| 15 | 50 | 1.000 | 2.8436e-10 | 0.3 | 1.5302e-11 | 2.6 | 25.8 | 63.7 |
| 25 | 50 | 1.000 | 2.8462e-10 | 0.2 | 1.4808e-11 | 2.1 | 27.6 | 61.6 |
| 40 | 40 | 1.000 | 2.8606e-10 | 0.2 | 1.4586e-11 | 2.1 | 28.5 | 60.7 |
| 60 | 68 | 1.000 | 2.8562e-10 | 0.2 | 1.4516e-11 | 1.3 | 28.5 | 60.4 |
| 80 | 52 | 1.000 | 2.8567e-10 | 0.2 | 1.4546e-11 | 1.4 | 29.7 | 60.5 |
| 100 | 40 | 1.000 | 2.8549e-10 | 0.2 | 1.4475e-11 | 1.2 | 28.1 | 60.2 |
| 150 | 26 | 1.000 | 2.8598e-10 | 0.1 | 1.4289e-11 | 1.2 | 27.9 | 59.4 |
| 200 | 30 | 1.000 | 2.8650e-10 | 0.1 | 1.4201e-11 | 1.1 | 27.8 | 59.1 |
| 300 | 24 | 1.000 | 2.8659e-10 | 0.1 | 1.4278e-11 | 1.1 | 28.3 | 59.4 |
| 500 | 12 | 1.000 | 2.8650e-10 | 0.1 | 1.4302e-11 | 1.2 | 29.0 | 59.5 |
| 1000 | 8 | 1.000 | 2.8591e-10 | 0.1 | 1.4306e-11 | 0.5 | 27.7 | 59.5 |

**Table S17  DNA damage yields per beam quality**

**folded over the spectrum of electrons the photons set in motion;**

**'no delta corr.' repeats the fold without removing delta rays**

| beam | <E_e> | Y_DSB | err% | DSB/Gy/cell | complex% | /Co-60 | no delta corr. |
|---|---|---|---|---|---|---|---|
| 40 kV | 25.1 | 1.5054e-11 | 1.5 | 62.6 | 27.3 | 1.051 | 1.041 |
| 50 kV | 27.0 | 1.5096e-11 | 1.3 | 62.8 | 27.4 | 1.054 | 1.043 |
| 60 kV | 28.2 | 1.5137e-11 | 1.2 | 63.0 | 27.4 | 1.057 | 1.046 |
| 80 kV | 29.5 | 1.5220e-11 | 1.1 | 63.3 | 27.4 | 1.063 | 1.050 |
| 100 kV | 29.9 | 1.5276e-11 | 1.0 | 63.6 | 27.3 | 1.067 | 1.053 |
| 120 kV | 30.3 | 1.5299e-11 | 0.9 | 63.6 | 27.3 | 1.069 | 1.054 |
| 200 kV | 31.5 | 1.5287e-11 | 0.9 | 63.6 | 27.3 | 1.068 | 1.053 |
| 100 kV +0.1Cu | 31.3 | 1.5342e-11 | 1.0 | 63.8 | 27.3 | 1.072 | 1.056 |
| 100 kV +0.2Cu | 31.8 | 1.5392e-11 | 0.9 | 64.0 | 27.3 | 1.075 | 1.059 |
| Co-60 | 725.0 | 1.4318e-11 | 0.5 | 59.6 | 28.1 | 1.000 | 1.000 |
